# The Chalcidoidea bush of life – a massive radiation blurred by mutational saturation

**DOI:** 10.1101/2022.09.11.507458

**Authors:** Astrid Cruaud, Jean-Yves Rasplus, Junxia Zhang, Roger Burks, Gérard Delvare, Lucian Fusu, Alex Gumovsky, John T. Huber, Petr Janšta, Mircea-Dan Mitroiu, John S. Noyes, Simon van Noort, Austin Baker, Julie Böhmová, Hannes Baur, Bonnie B. Blaimer, Seán G. Brady, Kristýna Bubeníková, Marguerite Chartois, Robert S. Copeland, Natalie Dale-Skey Papilloud, Ana Dal Molin, Chrysalyn Dominguez, Marco Gebiola, Emilio Guerrieri, Robert L. Kresslein, Lars Krogmann, Emily Moriarty Lemmon, Elizabeth A. Murray, Sabine Nidelet, José Luis Nieves-Aldrey, Ryan K. Perry, Ralph S. Peters, Andrew Polaszek, Laure Sauné, Javier Torréns, Serguei Triapitsyn, Ekaterina V. Tselikh, Matthew Yoder, Alan R. Lemmon, James B. Woolley, John M. Heraty

## Abstract

Capturing phylogenetic signal from a massive radiation can be daunting. The superfamily Chalcidoidea is an excellent example of a hyperdiverse group that has remained recalcitrant to phylogenetic resolution. Chalcidoidea are mostly parasitoid wasps that until now included 27 families, 87 subfamilies and as many as 500,000 estimated species. We combined 1007 exons obtained with Anchored Hybrid Enrichment with 1048 Ultra-Conserved Elements (UCEs) for 433 taxa including all extant families, over 95% of all subfamilies and 356 genera chosen to represent the vast diversity of the superfamily. Going back and forth between molecular results and our collective morphological and biological knowledge, we detected insidious bias driven by the saturation of nucleotide data and highlighted morphological convergences. Our final results are based on a concatenated analysis of the least saturated exons and UCE data sets (2054 loci, 284,106 sites). Our analyses support a sister relationship with Mymarommatoidea. Seven of the previously recognized families were not monophyletic, so foundations for a new classification are discussed. Biology appears potentially more informative than morphology, as illustrated by the elucidation of a clade of plant gall associates and a clade of taxa with planidial first-instar larvae. The phylogeny suggests a shift from smaller soft-bodied wasps to larger and more heavily sclerotized wasps. Deep divergences in Chalcidoidea coincide with an increase in insect families in the fossil record, and an early shift to phytophagy corresponds with the beginning of the “Angiosperm Terrestrial Revolution”. Our dating analyses suggest a Middle Jurassic origin of 174 Ma (167.3-180.5 Ma) and a crown age of 162.2 Ma (153.9–169.8 Ma) for Chalcidoidea. During the Cretaceous, Chalcidoidea underwent a rapid radiation in southern Gondwana with subsequent dispersals to the Northern Hemisphere. This scenario is discussed with regard to knowledge about host taxa of chalcid wasps, their fossil record, and Earth’s paleogeographic history.

## INTRODUCTION

With the increasing use of target-enrichment sequencing, much progress has been made in our understanding of the tree of life, but efforts remain unequal among taxonomic groups. Insects are arguably the most species-rich terrestrial organisms (Stork 2018), but so far only a few phylogenomic hypotheses using representative sampling have been published at the family level and above. Yet, in this “century of extinction”, filling the gap between biodiversity and our knowledge of it should be a top priority (Dubois 2003). In hyperdiverse groups, the scarcity of representative phylogenetic trees is primarily linked with the hyperdiversity itself. First, collaborative sampling efforts must be considerable to achieve as complete a representative sampling as possible, however the increasing burden of restrictive access to specimens and regulations that hinder sharing of specimens makes this task even more difficult (Prathapan et al. 2018). Second, there is a lack of well-trained taxonomists (Wheeler 2014; Britz et al. 2020; Engel et al. 2021) who are able to: 1) embrace the complexity of an immensely diverse group through identification or description of species; 2) assign fossils to groups and provide calibrations for dating analyses; or, 3) make a critical evaluation of the phylogenetic trees obtained from molecular data (Wiens 2004). Lastly, as new species and genera are constantly discovered, we are in dire need of comprehensive and reliable phylogenetic hypotheses that will all allow the accurate delimitation and placement of taxa that often suffer from a lack of scientific interest (Wägele et al. 2011). Yet, this world contains vast numbers of yet undiscovered taxa of tremendous ecological and economic importance (Foottit and Adler 2009).

While genomic data offer great promise to resolve the tree of life, analytical challenges must be overcome. When more markers are analyzed, the probability of observing conflicting signal among them increases (Kumar et al. 2012; Philippe et al. 2017; Zhang et al. 2020; Zhang et al. 2022). Highly supported trees can be inferred that, without feedback from taxonomists, are considered to be accurate even though inferences may be flawed (Wiens 2004; Zhang et al. 2022). Heterogeneity in base composition and in evolutionary rates inferred for both taxa and markers are major causes of analytical bias in phylogenomic analyses (e.g., Boussau et al. 2014; Romiguier and Roux 2017; Borowiec et al. 2019; Rasplus et al. 2021). Recent studies have brought to light an old nemesis, mutational saturation (Philippe and Forterre 1999), as a potentially major source of errors, especially for deep-time inferences (Borowiec 2019; Borowiec et al. 2015; Duchêne et al. 2022). Ideally, to detect and reduce inference biases, analyzed matrices should be rich both in taxa (Heath et al. 2008) and molecular characters of different origins [e.g., coding vs non-coding, Reddy et al. 2017]. They should also be mixed with or interpreted in the light of morphology biology and other data types (Wiens 2004), that can also be misleading and must be interpreted cautiously. Thorough analysis of very large data sets that implement many proof checking steps is computationally intensive. In addition, it is still impossible to perfectly describe evolutionary processes with mathematical models, which inevitably introduce bias (Kumar et al. 2012; Reddy et al. 2017). Hence, resolving the tree of life of ancient, poorly known and hyperdiverse groups requires determination and humility. When independent data types do not converge towards the same results, molecular trees should certainly be acknowledged as valuable contributions, but considered only as hypotheses, instead of being hailed as “the resolved tree of life”.

Chalcidoidea (jewel wasps, chalcidoid wasps, hereafter called chalcid wasps) are among the most species rich, ecologically important, biologically diverse and morphologically disparate groups of terrestrial organisms. These minute wasps (mostly 0.5–2 mm in size) are numerically abundant and ubiquitous in almost every terrestrial habitat on Earth. Their diversity is staggering, with an estimated 500,000 species placed in 27 families (among which two are extinct) and 87 subfamilies (among which 3 are extinct) and several *incertae sedis* taxa (Table S1a). 2,731 genera and 27,021 species have been described so far [Universal Chalcidoidea Database (Noyes 2019) currently being migrated to TaxonWorks (TaxonWorks Community, 2022). Although they are mostly parasitoids, phytophagous species are known from nine families (Böhmová et al. 2022). Their animal host range includes 13 insect orders, spiders, ticks, mites, pseudoscorpions and even gall-forming nematodes (Austin et al. 1998; Gibson et al. 1999). Chalcid wasps attack all life stages of their hosts from eggs to adults, as internal or external parasitoids. They can be primary, secondary, or even tertiary parasitoids, and some large lineages are characterized by male and female larvae developing differently on the same host, or more commonly on different hosts [heteronomy; Hunter et al. (2001)]. In the most extreme form of heteronomy, males can be obligate hyperparasitoids of females of their own species. The economic importance of Chalcidoidea in pest management is unparalleled and they are widely used in biological control programs against major pests throughout the world (Noyes and Hayat 1994; Heraty 2009). Only recently have we begun to attempt an understanding of their phylogenetic relationships (Munro et al. 2011; Heraty et al. 2013; Peters et al. 2018; Zhang et al. 2020), but our progress is still very incomplete.

A few studies have addressed higher-level relationships within Chalcidoidea, although with only a sparse sampling of genes (Sanger data sets) or taxa (NGS data sets). Munro et al. (2011) used 18S+28S ribosomal DNA for 649 species of Chalcidoidea in 19 families and 343 genera; Heraty et al. (2013) used a combination of 18S+28S+morphology for 283 species in 19 families and 268 genera; Peters et al. (2018) analyzed 3,239 genes from transcriptomes for 37 species in 16 families and 35 genera; and Zhang et al. (2020) extended the transcriptome data set to 5,591 genes for 55 species in 17 families and 48 genera. Despite these efforts, the higher-level relationships of Chalcidoidea still remain largely unresolved. Genome-scale data (transcriptomes) have proven particularly frustrating, presumably because of the lack of signal associated with an old, rapid radiation and/or of the increasing probability of observing conflicting signals between markers (Peters et al. 2018; Zhang et al. 2020). When chalcid wasps are included in studies of the Hymenopteran tree of life, conflicts or lack of signal that are reflected in poor statistical support of (some) nodes are highlighted in all data sets: Branstetter et al. (2017) 854 Ultra-Conserved Elements (UCEs), 9 species of Chalcidoidea in 9 families and 9 genera; Peters et al. (2017) 3,256 protein coding genes, 6 species in 6 families and 6 genera; Tang et al. (2019) mitochondrial genomes, 7 species in 6 families and 7 genera. In addition, the tree inferred from mitochondrial genomes cannot be reconciled with those inferred from transcriptomes or UCEs. Finally, with the scarce taxonomic sampling of previously published phylogenomic data sets, it has not been possible to test the monophyly of chalcid families or subfamilies that were questioned by Sanger data sets and morphology (e.g., Aphelinidae, Pteromalidae; (Munro et al. 2011; Heraty et al. 2013), or to resolve the position of *incertae sedis* taxa that may represent independent, possibly old, evolutionary lineages. More importantly, reduced taxonomic sampling coupled with poor resolution of phylogenetic trees limits our understanding of the drivers that fueled this outstanding diversity of life forms and biologies across space and time.

This overview of previous works suggests that, although the early evolution of Chalcidoidea likely represents a difficult phylogenetic problem, we may advance our knowledge of their tree of life through the acquisition and careful comparative and combined analysis of different types of molecular markers obtained from a representative set of species. Unsurprisingly, given their extreme morphological disparity and rampant convergence, a major collaborative effort to provide a resolved morphological tree for the superfamily largely failed (233 morphological characters scored on 283 species in 19 families; Heraty et al. 2013), although several family-level groups were recovered that were not found in the analyses of ribosomal markers. Within Chalcidoidea, when phylogenomic studies focused on smaller taxonomic units, the results were either in strong agreement with morphology, behavior or biogeographic hypotheses (e.g., Baker et al. 2020; Rasplus et al. 2020), or strongly conflicting on some areas of the tree with intuitive and previously supported hypotheses (e.g., Cruaud et al. 2021; Zhang et al. 2022). When there is conflict, there is always the possibility that properties of the genomic data or confounding signal in morphological/biological data may be affecting the results. But only with a thorough analysis and vetting of the data can we begin to either understand the issues and potential for systematic bias or morphological convergence.

In this study, we brought together taxonomists and museum curators to assemble a taxon- and marker-rich data set (exons + UCEs and their flanking regions) for Chalcidoidea. To find our way through a forest of phylogenetic trees, we evaluated topologies obtained from each genomic data type in the light of our morphological/biological knowledge. Taxa/groups for which we inferred unlikely relationships were used to detect and reduce potential bias in the molecular data sets. Conversely, morphological/biological data were re-examined to assess (hidden) support for unexpected relationships (reciprocal illumination, Hennig 1966; Mooi and Gill 2016). We use the combined exons+UCEs least biased data set to provide the foundation for a new classification of the superfamily that will be published in another manuscript (Burks et al. submitted), discuss its evolutionary history, and infer a timeline for its origin and worldwide colonization.

## MATERIALS AND METHODS

### Taxonomic sampling

Representatives of all extant families, 80 extant subfamilies (95.2%), 68 extant tribes (77.3%) and 356 genera (13%) of chalcid wasps were included. Representatives of 8 *incertae sedis* taxa at the suprafamilial or tribal levels and of one new subfamily were also included. A total of 433 taxa (414 ingroups and 19 outgroups) were analyzed (Table S1b), of which, 414 had sequences for exons while 407 had sequences for UCEs. Exons and UCEs were obtained from the same species in 57% of the ingroup taxa, while congeneric specimens were used in the remaining 43%. For seven taxa (4 outgroups, 3 ingroups), exons and UCEs were obtained from species in different genera (all very closely related for ingroups). Our outgroups include a diverse array of Proctotrupomorpha, including Platygastroidea (2 genera), Cynipoidea (5 genera), Proctotrupoidea (2 genera), Diaprioidea (5 genera), and Mymarommatoidea (2 genera). These outgroups form a paraphyletic grade to Chalcidoidea in all recent analyses of Hymenoptera relationships (Heraty et al. 2011; Klopfstein et al. 2013; Branstetter et al. 2017; Peters et al. 2017). The supplementary documents also include results obtained with a larger exon data set of 520 taxa (494 ingroups and 26 outgroups, hereafter referred to as the AHE520 data set) from which the 414 taxa to be paired with UCEs (hereafter referred to as the AHE414 data set) were extracted (Table S1b). As compared to the AHE414 data set, sampling within some families was increased in the AHE520 data set but the same evolutionary lineages were included. We chose not to combine all of the taxa in the AHE520 data set with the UCE data set to 1) avoid potential issues with missing data, 2) enable better comparison between properties and phylogenetic signal brought by the exons and UCEs, and 3) decrease computational burden. Nevertheless, trees obtained with the AHE520 data set (exons with RY coding of the third codon position and coded as amino acids) were compared with those obtained with the AHE414, the UCE and the combined data sets.

### Library preparation and sequencing

#### Exons

Exons were obtained for 520 taxa following the protocol in Zhang et al. (2022), of which 51 taxa were retrieved from previously published transcriptomes or genomes (Peters et al. 2018; Zhang et al. 2020), and 469 taxa (363 used in combined dataset) were enriched using the anchored hybrid enrichment (AHE) probe sets (*Hym_Ich set* or *Hym_Cha set*). See supplemental methods for details on library preparation and sequencing.

#### UCEs

UCEs for 116 taxa were retrieved from previous studies (Cruaud et al. 2019; Rasplus et al. 2020; Rodriguez et al. 2021). For the remaining 291 taxa, library preparation followed Cruaud et al. (2019). Specimens were enriched in 1432 UCEs using the 2749 probes designed by Faircloth et al. (2015) (myBaits UCE Hymenoptera 1.5Kv1 kit; Arbor Biosciences). See supplemental methods for details on library preparation and sequencing.

### Assembly of data sets

#### Exons

Assembly of loci, orthology assessment and contamination checking for the AHE520 data set followed Zhang et al. (2022) (see also supplementary methods). Exons were selected of 414 taxa that were compatible for data combination with the UCE taxa, either as the same extraction, same species, same genus, or in 3 cases as closely related genera (Table S1b). However, the assembly protocol used in Zhang et al. (2022) was overly stringent, resulting in 28.5% of missing or ambiguous nucleotide calls. Therefore, raw sequence data for these 414 taxa were re-processed with HybPiper to increase completeness (Johnson et al. 2016). All exons of the AHE520 data set were used as a reference database (final % of ambiguous/missing nucleotides = 19.1%; see supplementary methods for details).

#### UCEs

Raw data cleaning and assembly into loci followed Cruaud et al. (2019). Only UCEs that had a sequence for at least 50% of the samples were retained for analysis (N=1048). More details on the assembly of data sets can be found in the supplementary methods.

### Quality controls of sequence data

From this point, methods only refer to the AHE414 data set that was formally compared and then combined with the UCE data set and to the UCE data set. All details for the methods used for the AHE520 data set can be found in the supplementary methods.

#### Alignment cleaning

Loci (exons and UCEs) were aligned with MAFFT using the –linsi option (Katoh and Standley 2013). Exons were translated to amino acids using EMBOSS (Rice et al. 2000) and sequences with stop codons were removed. Two successive rounds of TreeShrink (Mai and Mirarab 2018) were performed on each locus (exons analyzed as nucleotides) to detect and remove abnormally long branches in individual gene trees. The per-species mode was used and b (the percentage of tree diameter increasing from which a terminal should be removed) was set to 20. Loci were re-aligned with MAFFT after each round of TreeShrink. Gene trees were inferred with IQ-TREE v 2.0.6 (Nguyen et al. 2015; Minh et al. 2020) with the best fit model selected by ModelFinder (Kalyaanamoorthy et al. 2017). Positions with > 50% gaps and sequences with > 25% gaps were removed from the alignments of UCEs using SEQTOOLS (package PASTA; Mirarab et al. 2014) to speed up inference of individual gene trees.

#### Contaminations

A BLAST search of all DNA sequences on themselves (i.e., by using sequences of all exons/UCEs for all samples as query and target sequences) was performed (blastn with -evalue 1e-20 and -max_target_seqs 2). Only hits for which the same locus was identified as both target and query sequence and for which samples were different for target and query sequences were kept for downstream analysis. Putative contaminations were identified using a script that scored hits according to four criteria appropriate for our data (cf. https://github.com/mjy/cgq for details): 1) *Taxon Difference*: either subfamily or family was different between target and query sequences; 2) *Proportional Difference*: the percentage of similarity between target and query was > 99.95; 3) *Proportional Length Difference*: length of match divided by the smaller of length of the target and query was > 0.95; 4) *Plate Similarity*: target and query sequences were obtained from specimens processed on the same plate for DNA extraction/library preparation. When a criterion was met, a score of 1 was attributed, otherwise the score was 0. A composite score was calculated as the sum of criteria 1 to 4. When the composite score equaled 4 both target and query loci were considered as potentially contaminated. The ratio of query species DNA concentration to target species DNA concentration was calculated. When ratio < 0.3 the sequence with the smaller qubit concentration was excluded from the data set, otherwise both sequences were excluded.

### Properties of taxa, loci, trees and exploration of bias

#### Workflow to detect and decrease bias

Due to constraints imposed by the computational resources required to analyze our large data set, we adopted a pragmatic approach to detect possible sources of bias in the data. We focused on only two properties of loci that were, according to the literature, the most likely to impact our inferences: saturation (Borowiec et al. 2015; Borowiec 2019; Duchêne et al. 2022) and GC content (Bossert et al. 2017; Cruaud et al. 2021). First, we analyzed data subsets that were less and less saturated and analyzed whether resulting topologies had a better fit to morphological and/or biological data and/or previously published hypotheses. Reciprocally, morphological/biological data were re-examined to highlight convergences and assess support for unexpected relationships (Mooi and Gill 2016). To assess fit to morphological data we determined whether several currently recognized family-level taxa were monophyletic in resulting trees. To assess fit to biological data, we determined whether two clades supported by biological properties (e.g., a clade of plant gall associates and a clade of taxa with planidial first-instar larvae) were monophyletic in resulting trees. To incrementally decrease saturation, exons were analyzed as nucleotide sequences, with RY coding of the third codon position of each amino acid (Delsuc et al. 2003) (script RYplace.py; Ballesteros and Hormiga 2016) and as amino acids. Three corresponding exon data sets were built for the AHE414 data set, hereafter referred to as exons, exonsRY and exonsAA (Table 1). Saturation in UCEs was reduced by filtering out gappy positions with three different thresholds until we reached a level of saturation for the concatenated UCE data set that was comparable to that of the concatenated exonsAA data set (Figure 1). Nucleotide positions in each aligned UCE were kept only when they were present in at least 50%, 70% or 90% of the taxa. In addition, to avoid bias due to misalignment of extremities of short sequences, sequences with more than 25% gaps were removed from each UCE. Removal of gappy positions and gappy sequences was performed with SEQTOOLS. Three corresponding UCE data sets were thus built, hereafter referred to as UCEs50-25, UCEs70-25 and UCEs90-25 (Table 1). Meanwhile, we explored whether other properties of loci (GC content and heterogeneous evolutionary rates) could explain placements of taxa that were unexpected in exon and UCE topologies based on morphological and/or biological data. The least saturated exon and UCE data sets turned out to have the better fit to both morphology and biology. Consequently, we tested whether or not removing 5% of the most saturated exons and 5% of the most GC-biased UCEs could further improve the results. However, it was not possible to assess whether this last attempt reduced bias or simply resulted in loss of phylogenetic signal. Given that the exonsAA and UCEs90-25 data sets produced results that were most congruent with both morphological and biological data (Figure 2), they were considered as the best hypotheses.

**Figure 1.**
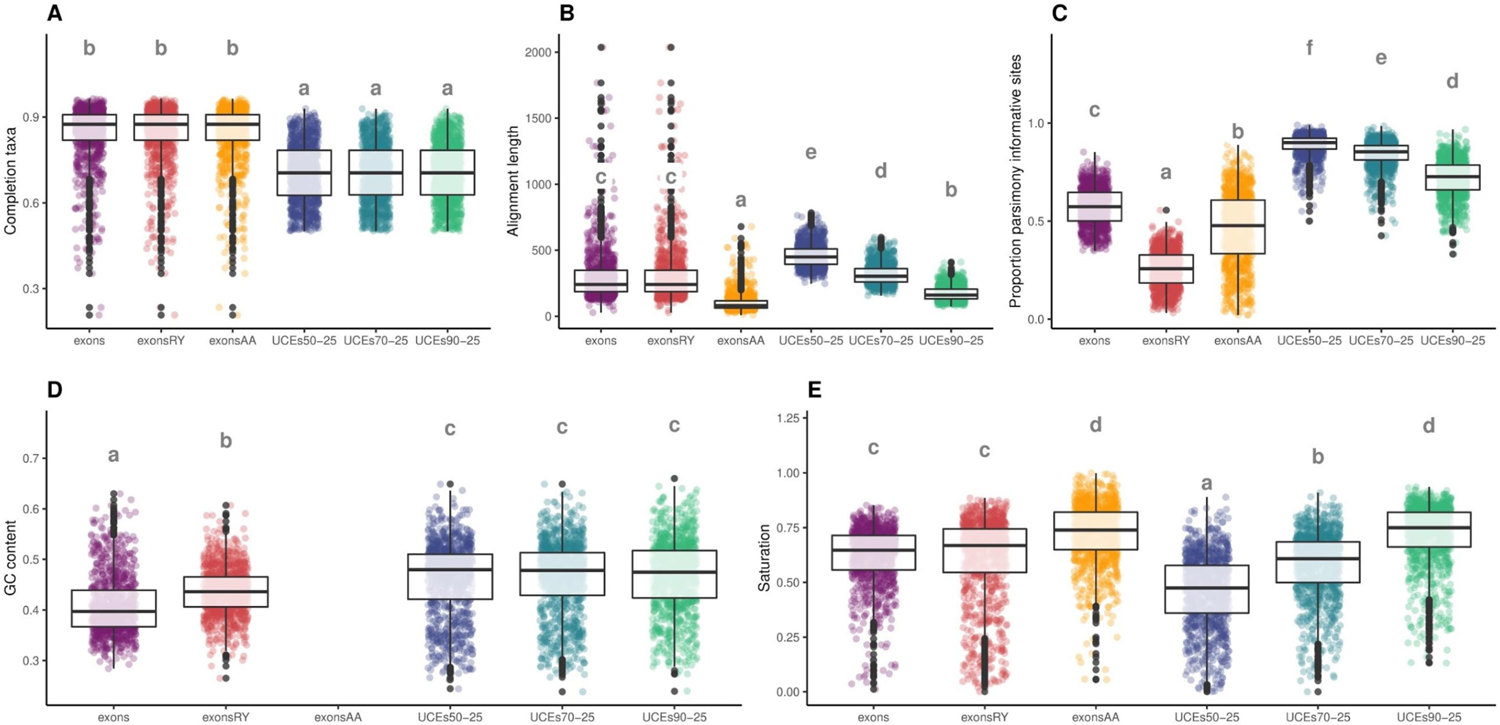
Comparison of properties of the analyzed data sets. Data sets (AHE414 and UCEs) are described in Table 1. For each panel, letters above box plots reflect pairwise comparisons of marginal means estimated from the best fit models; distributions sharing a letter do not differ significantly. Points: raw data (Table S2a). Saturation was assessed by calculating the R squared of the linear regression of uncorrected p-distances against inferred distances in individual gene trees.

**Figure 2.**
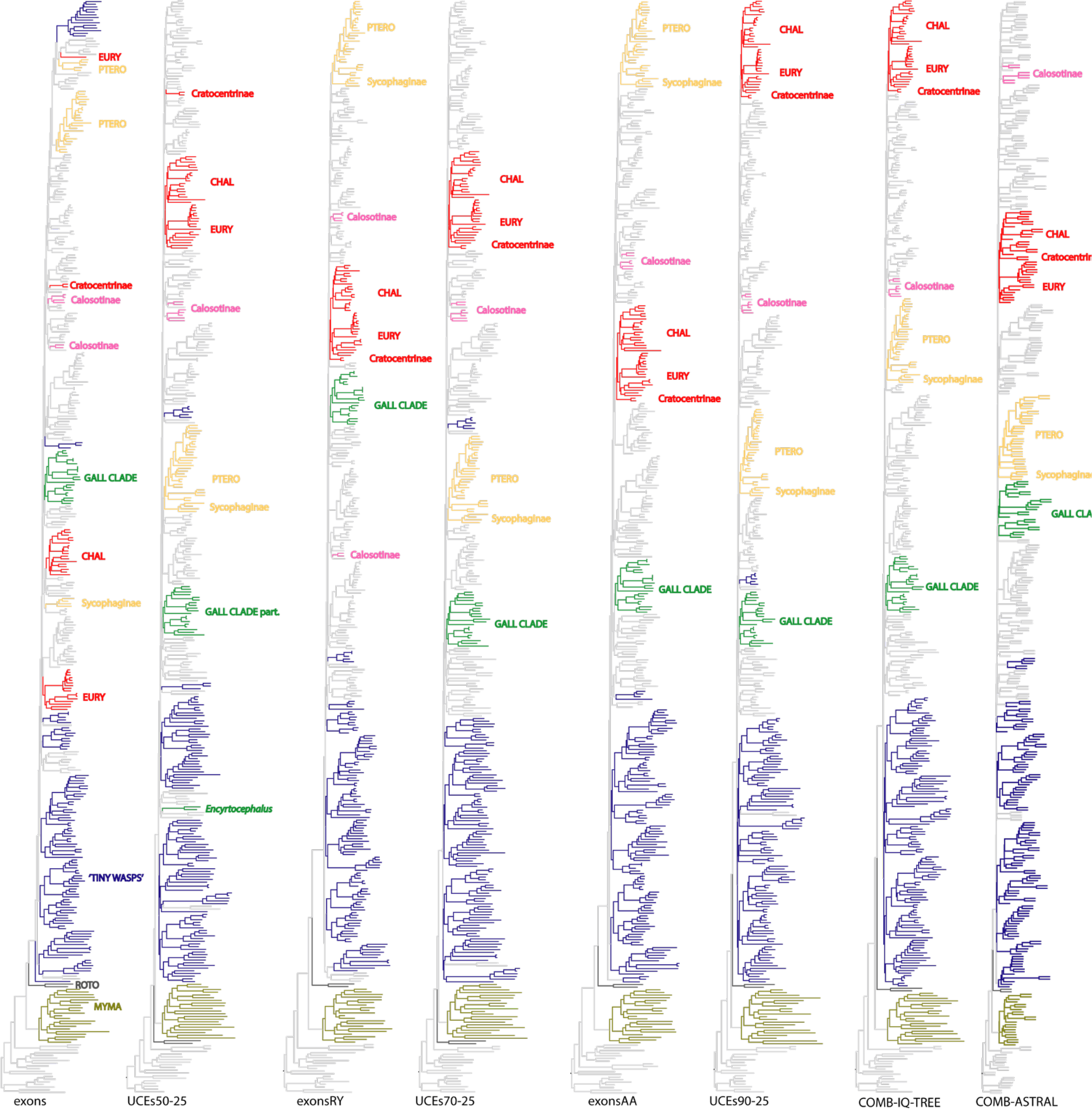
Overview of the topologies obtained with the different data sets. Data sets (AHE414 and UCEs) are described in Table 1 and trees are available in Figure S1 and Appendix S2. Groups that are discussed in text are highlighted. Only IQ-TREE trees are shown for the exons and UCE trees (unpartitioned data sets) but both IQ-TREE (data set partitioned by type of markers exonsAA vs UCEs90-25) and ASTRAL trees are shown for the combined data set. ROTO=Rotoitidae; CHAL=Chalcididae; EURY=Eurytomidae; GALL = gall clade (see text); MYMA=Mymaridae; PTERO = group of Pteromalid wasps (Austroterobiinae; part Colotrechninae; Miscogastrinae; part Ormocerinae; Otitesellinae; Pteromalinae; Sycoecinae; Sycoryctinae); “Tiny Wasp clade” (see text).

**Table 1.**
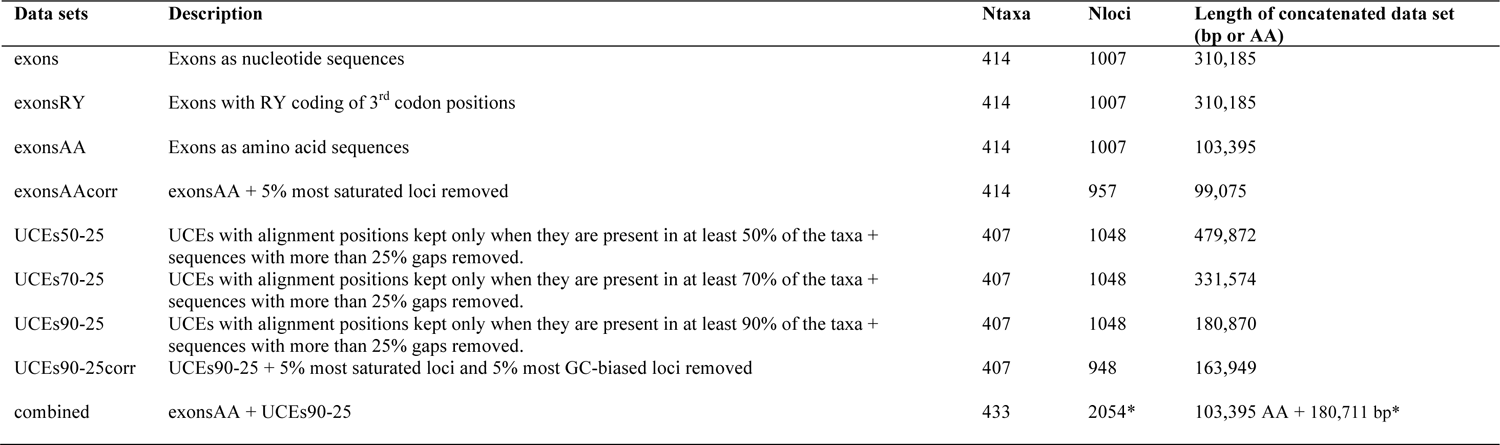
Description of the exons (AHE414) and UCE data sets. Detailed properties are given in TableS2b. *Note that the locus shared among the exons and the UCE data set was removed from the UCE data set before running the combined analysis.

#### Calculation of properties

GC content of taxa was calculated with AMAS (Borowiec 2016). Long branch (LB) score heterogeneity for taxa in trees (taxon’s percentage deviation from the average pairwise distance between taxa on a given tree) was used as a proxy of evolutionary rate of taxa and was calculated with TreSpEx (Struck 2014). Properties of locus and concatenated data sets (length, proportion of parsimony informative sites; GC content; etc. Table S2a) were calculated with AMAS. Saturation of loci (R squared of the linear regression of uncorrected p-distances against inferred distances in individual gene trees) was calculated as in Borowiec et al. (2015).

#### Statistical analyses

Analyses were performed in R (R Core Team 2018). Hierarchical clustering of taxa based on GC content and LB scores was performed with the package *cluster* (Maechler et al. 2018). Strength and direction of association between variables were assessed 1) with Spearman’s rank correlation using *PerformanceAnalytics* (Peterson and Carl 2018) or 2) by fitting linear models (with log-transformation of variables when relevant (Ives 2015). Significant deviations from model assumptions (normality of residuals, homoscedasticity) and absence of highly influential data points were detected with *DHARMa* (Hartig 2022) and *performance* (Lüdecke et al. 2021). A likelihood ratio test was used to test the significance of fixed factors. A Tukey post-hoc test was used when more than 2 groups were compared (packages *emmeans* (Lenth 2021) and *multcomp* (Hothorn et al. 2008)). Graphs were generated with *ggplot2* (Wickham 2016).

### Combined data set

To reveal potential hidden support, the exonsAA and UCEs90-25 data sets were combined (Table 1). Before combination, overlap between exons and UCEs was tested with reciprocal BLAST (UCEs not trimmed with SEQTOOLS; blastn with -evalue 1e-20). Only a single locus was shared between data sets and it was removed from the UCE data set before combination.

### Phylogenetic inference

Data sets were analyzed with concatenation (using IQ-TREE 2.0.6) and tree reconciliation (using ASTRAL-III, Zhang et al. 2018) approaches. For the concatenation approach, loci were merged and the resulting data set was analyzed 1) without partitioning, 2) with one partition for each locus and 3) with one partition for each data type (combined data set only; 1 partition for the exons another for the UCEs). Best fit models were selected with the Bayesian Information Criterion (BIC) as implemented in ModelFinder. FreeRate models with up to ten categories of rates were included in tests for the unpartitioned exon and UCE data sets, but only common substitution models were tested when data sets were partitioned by locus. The candidate tree set for all tree searches was composed of 98 parsimony trees + 1 BIONJ tree and only the 20 best initial trees were retained for NNI search. Statistical support of nodes was assessed with ultrafast bootstrap (UFBoot) (Minh et al. 2013) with a minimum correlation coefficient set to 0.99 and 1,000 replicates of SH-aLRT tests (Guindon et al. 2010). Gene (gCF) and site (sCF) concordance factors (Minh et al. 2020) were also calculated in IQ-TREE.

For ASTRAL analyses, nodes in gene trees with UFBoot support lower than 90 were collapsed (perl script AfterPhylo.pl, Zhu 2014) before reconciliation. Statistical support of nodes was assessed with local posterior probabilities (local PP) as implemented in ASTRAL-III. Distance between a node and its parent node was calculated with the R package ape (Paradis and Schliep 2018). RF distances (Robinson and Foulds 1981) among recovered trees were calculated with RAxML-NG_v0.9.0 (Kozlov et al. 2019).

### Divergence time estimates

Time calibrated trees were generated with MCMCtree (Yang and Rannala 2006). Twenty-one fossils were used as calibration priors and uniform distributions were used as calibration densities (Table S3a; Appendix S1). Analyses were run with uncorrelated relaxed clock models. The combined IQ-TREE tree (partitioning by data type) was used as the input tree. Five data sets, each composed of 10,000 amino acid sites randomly selected (custom script in Rougerie et al. 2022) from the exonsAA partition + 10,000 nucleotide sites randomly selected from the UCEs90-25 partition, were used as sequence data to make computation tractable. Each data set was partitioned into two partitions exonsAA (WAG+G model) and UCEs (GTR+G model). Four chains were run for each data set; 20,000 generations were discarded as burnin and chains were run for 2M generation with sampling every 200 generations. Convergence was assessed in Tracer (Rambaut et al. 2018). Possible conflicts between priors and data were assessed by running MCMCtree without sequence data. Posterior estimates obtained with the different data sets were compared and combined with LogCombiner 2.6.0 (Bouckaert et al. 2019).

### Historical biogeography

Distributions of species in each genus were mined from the Universal Chalcidoidea Database (Noyes 2019). Occurrences were double checked by experts of each taxonomic group and modified if needed. Species used as biocontrol agents or accidentally introduced with their host (plant or insect) (Rasplus et al. 2010; Noyes 2019) were removed from the list before compiling genus-level distribution data. Genera were scored as present/absent in the six following biogeographical areas: Neotropical, Nearctic, Afrotropical, Palaearctic, Oriental, Australasian (Table S4a). Any genus for which a single species occurs at the boundary of the transition zone between two areas while all other species occur in only one area was coded as present only in the latter area. Ancestral area estimations were performed using the R package *BioGeoBEARS 1.1.1* (Matzke 2014). The chronogram built with MCMCtree was used as input, but only one specimen per genus was included and outgroups were pruned to avoid artefacts. Dispersal-Extinction-Cladogenesis (DEC; Ree and Smith 2008), BAYAREALIKE (Landis et al. 2013) and DIVALIKE (Ronquist 1997) models were used with and without considering the jump parameter for founder events (+*J*; Matzke 2014). Model selection was performed based on statistical (AICc; Matzke 2021) and non-statistical (i.e., biological and geographical) considerations (Ree and Sanmartin 2018). The maximum number of areas that a species could occupy was set to 6. To consider the main geological events that occurred during the diversification of Chalcidoidea, we defined five time periods with different dispersal rate scalers: 1) from their mean crown age to 145 Ma (Jurassic); 2) 145 to 100 Ma (Early Cretaceous); 3) 100 to 66 Ma (Late Cretaceous); 4) 66 to 23 Ma (Paleogene); 5) 23 Ma to present (Table S4b).

## RESULTS

### Detection and reduction of inference bias

The initial set of exons (AHE414; Table 1) had a high number of loci recovered across taxa and was less saturated and less GC rich than the initial set of UCEs (UCEs50-25) that, in comparison, contained longer loci and more parsimony informative sites (Figure 1, Table S2a). Six groups of chalcid wasps (Sycophaginae, Cratocentrinae, Calosotinae, Mymaridae, Rotoitidae, *Encyrtocephalus*) were recovered in mostly unexpected positions in the trees inferred from the concatenation (IQ-TREE) or the reconciliation (ASTRAL) of these initial sets of markers (Figures 2, S1; Appendix S2). Sycophaginae (Agaonidae) either clustered with (UCEs50-25) or away from (exons) a group of pteromalids (Austroterobiinae; part Colotrechninae; Miscogastrinae; part Ormocerinae; Otitesellinae; Pteromalinae; Sycoecinae; Sycoryctinae). Cratocentrinae (Chalcididae) never clustered with other Chalcididae.

Calosotinae (Eupelmidae) was never recovered as monophyletic. Either Mymaridae (exons and ASTRAL UCEs50-25) or Rotoitidae (IQ-TREE UCEs50-25) were recovered as sister to all other chalcid wasps. *Encyrtocephalus* (Pteromalidae) was either recovered outside (IQ-TREE and ASTRAL UCEs50-25; ASTRAL exons) or inside of a clade of gall associates (IQTREE exons). As saturation decreased (from exons to exonsAA and from UCEs50-25 to UCEs90-25; Figure 1), placement of these taxa tended to become more and more similar between trees and concordant with current morphological hypotheses (Figures 2, S1). Clustering analyses of taxa properties (GC content and LB scores) did not provide evidence to consider that these unexpected placements were driven by compositional bias or long branch attraction (Figure S2; Table S5). Indeed, monophyletic groups with different GC content as well as polyphyletic groups with similar LB scores were recovered in the different trees. Interestingly, with decreasing saturation, groupings of taxa that were not expected based on morphological features but that are concordant with biological data appeared (i.e., a group clustering gall associates hereafter called “Gall clade” and a group of “Tiny wasps” hereafter called “Tiny Wasp clade” Figures 2, S1). Topological changes that may be attributed to reduction of mutational saturation are listed in Table S6.

Visual comparison of trees and RF distances (Figures 2, S1; Table S2c) showed that the exonsAA and UCEs90-25 data sets (i.e., the least saturated) produced IQ-TREE and ASTRAL trees that were the most similar, and that mostly agreed with morphological or biological expectations and prior taxonomic classifications. Further attempts in reducing bias by removing the most saturated and GC biased loci (exonsAAcorr and UCEs90-25corr data sets; Table 1) resulted in only a few topological changes in weakly supported regions of the topology (Figure S1). Therefore, to preserve possible hidden signal, we combined the exonsAA and UCEs90-25 data sets.

### Phylogenetic relationships

Readers can refer to Table S6 and Figure S3 for a detailed comparison of the exonsAA (414 taxa), UCEs90-25 and combined trees. Figure S3 also shows the AHE520AA tree (increased taxonomic sampling with exons coded as AA) for purpose of comparison. Notably, most higher-level clades (Figure S3, collapsed clades) are supported in all of the analyses, whereas the relationships between clades (Figure S3, vertical bars) are more variable but generally supported in all of the phylogenomic analyses.

More UCEs than exons supported the combined tree (gCF; Figure 3B), but a significant percentage of discordance was due to the lack of resolution of gene trees (gDFP). Distributions of sCF for nodes were identical for the two types of markers (Figure 3B). Statistical support was higher for longer branches (Figure S4A-B). Absolute RF distances between the exonsAA and the combined trees or the UCEs50-25 and the combined trees were close (10 more branches shared between the UCEs90-25 and the combined trees; Table S2c). The combined tree was thus considered as an acceptable compromise between the two types of markers. Differences were observed when the combined data set was partitioned by data type (AA vs UCEs) or by locus (Figure S1), but they were all observed in poorly supported sections of the topology. Given that short loci generated numerical instability for the estimation of model parameters, we favored a partitioning by type of markers.

**Figure 3.**
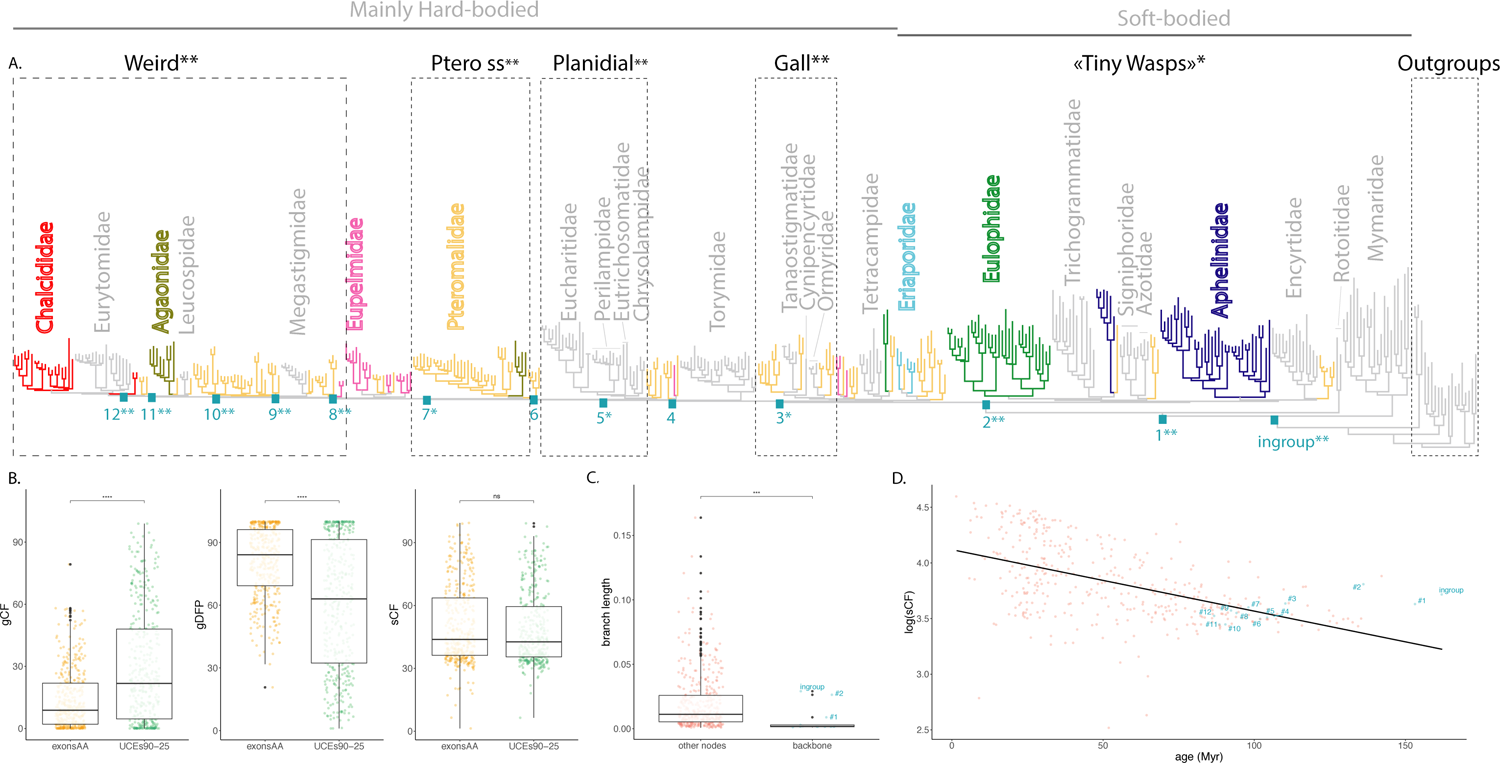
The Chalcidoidea bush of life. **A.** IQ-TREE tree obtained from the combined exonsAA+UCEs90-25 data sets (see also Figure S1). Monophyletic families are in grey, para- or polyphyletic families are in color. Higher level groups/clades discussed in text are highlighted with boxes. Statistical support for backbone nodes are shown with single (SH-aLRT ≥ 80% or UFboot ≥ 95%) or double stars (SH-aLRT > 80% and UFboot ≥ 95%) **B.** Contribution of the exonsAA and UCEs90-25 data sets to the combined tree. Gene concordance factor (gCF); gene discordance factor due to polyphyly (gDFP); site concordance factor averaged over 100 quartets (sCF). Points: raw data (Table S2d) **C.** Comparison of branch length for the backbone nodes and other ingroup nodes. Points: raw data (Table S2c). For panels B and C, letters above box plots reflect pairwise comparisons of marginal means estimated from the best fit models; distributions sharing a letter do not differ significantly. **D.** Correlation between node age and sCF (outgroups excluded). Points: raw data (Table S2e); line: regression curve for the best fit model (log linear model; P< 2.2e-16).

We observed that, for the deeper nodes, clades were inferred by IQ-TREE where ASTRAL inferred grades of lineages (Figures 2, S1), possibly as a result of hidden support for the first approach and uninformative gene trees for the second. Chalcidoidea were always recovered as monophyletic with strong support (Figures 3A, S1, S3). Of the 25 extant families (Table S1a), 18 were recovered as monophyletic with strong support while 7 were recovered as paraphyletic or polyphyletic (Figures 3A, S1, S3). The worst case, were the Pteromalidae, which were spread across the entire tree. Aphelinidae and Eulophidae were polyphyletic because of one genus in each family clustering away from the others: *Cales* and *Trisecodes,* respectively. Finally, Agaonidae (2 lineages), Chalcididae (2 lineages) and Eupelmidae (5 lineages) were polyphyletic, though Chalcididae was recovered as monophyletic in the ASTRAL results.

Some higher-level relationships were inferred that reflect biology more than morphology (Figure 3A). A clade of gall-associated wasps: Cynipencyrtidae + Ormyridae + Tanaostigmatidae + lineages of Pteromalidae (Epichrysomallinae + Melanosomellini); a clade of “Tiny Wasps” mostly associated with Hemiptera: Aphelinidae + Azotidae + Encyrtidae + Eulophidae (excluding *Trisecodes*) + Signiphoridae + Trichogrammatidae + certain lineages of Pteromalidae (Eunotini + *Idioporus* + Neodiparinae + Elatoidinae), and the planidial-larva clade: Eucharitidae + Perilampidae + Eutrichosomatidae + Chrysolampidae were recovered in the combined tree. One higher level grouping (hereafter referred to as the “Weird clade”) was unexpected based on morphology or biology.

All but two nodes (#1 and #2; Figure 3A) in the backbone were closer to each other than most ingroup nodes, suggesting near simultaneous old divergences for most nodes (Figure 3C). Only these two backbone nodes were recovered in the set of gene trees. Sixty-percent of the backbone nodes have SHaLRT and UFBoot higher than the suggested cut-off for validity (> 80% and > 95% respectively; IQ-TREE manual; Figures 3A-4, S1). A generic cut-off for sCF across a tree that spans such a long time makes little sense. Hence, we used the lowest sCF obtained for families that are well defined morphologically and biologically (Heraty et al. 2013): 34.3 for Trichogrammatidae. Using this value as a cut-off, 50% of the backbone nodes are supported (Figures 3A-4, S1). We observed a significant overall decrease in all statistical supports for nodes (gCF, sCF, UFBoot, SH-aLRT) with decreasing age (Figures 3D, S4C).

**Figure 4.**
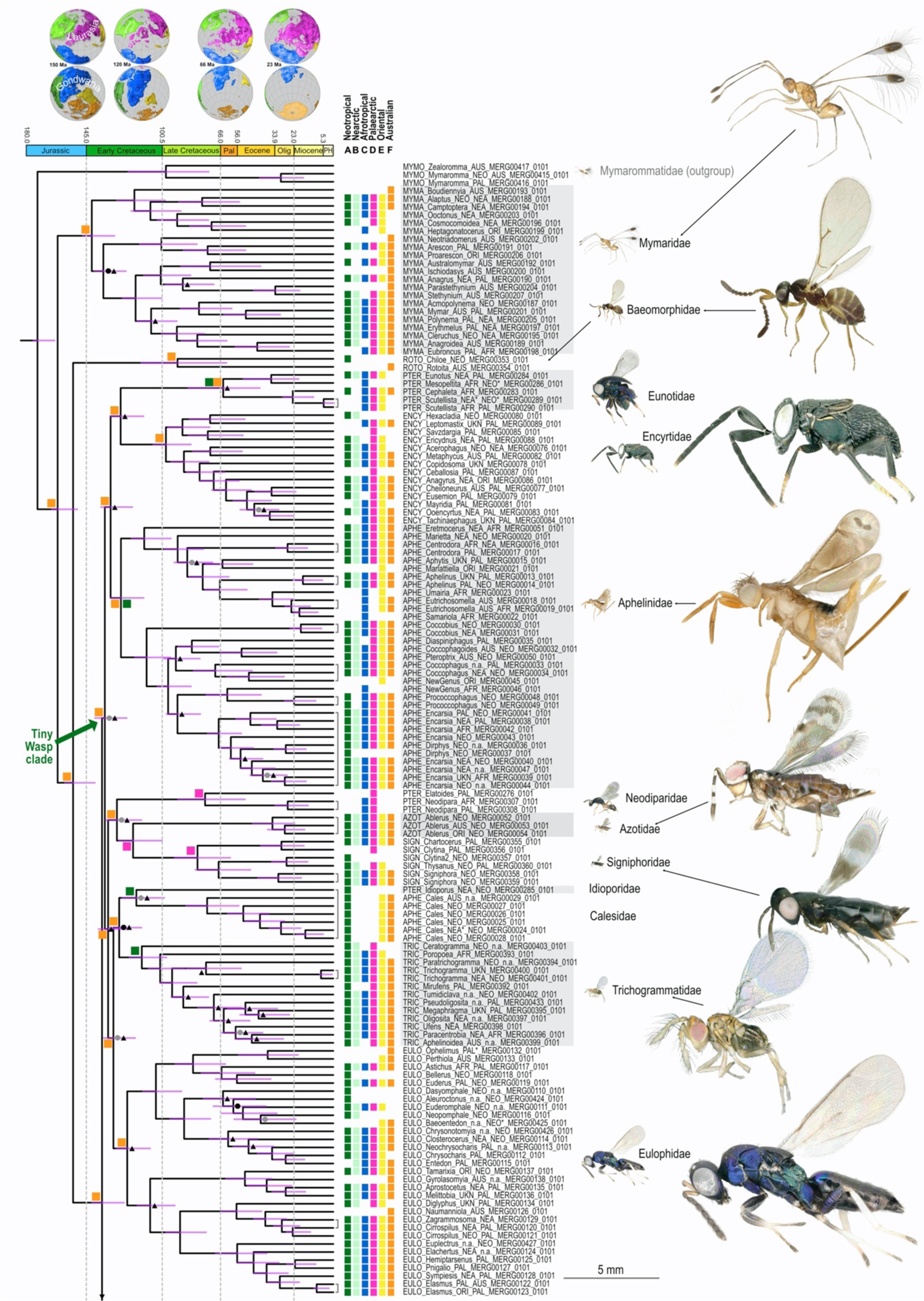
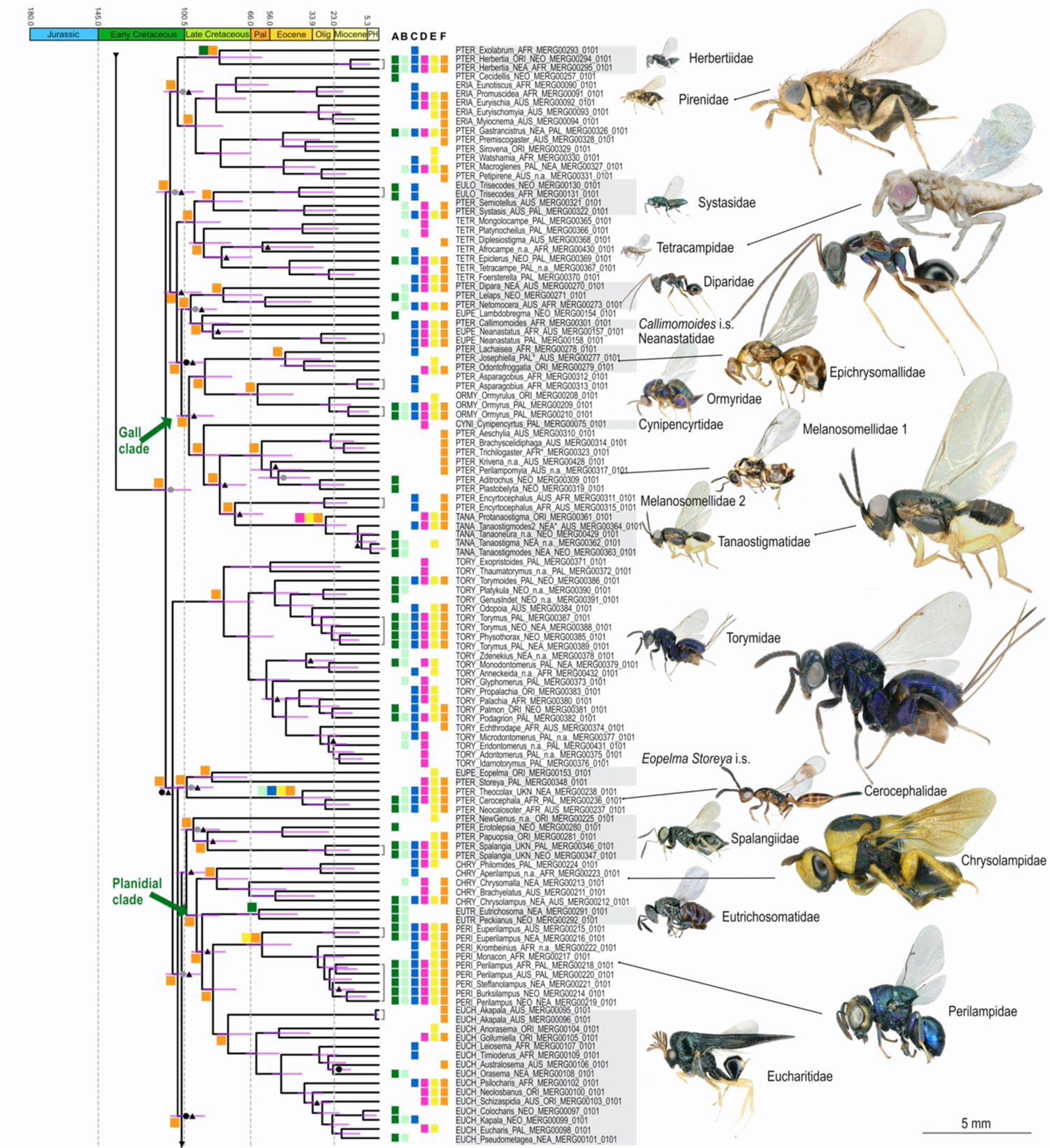
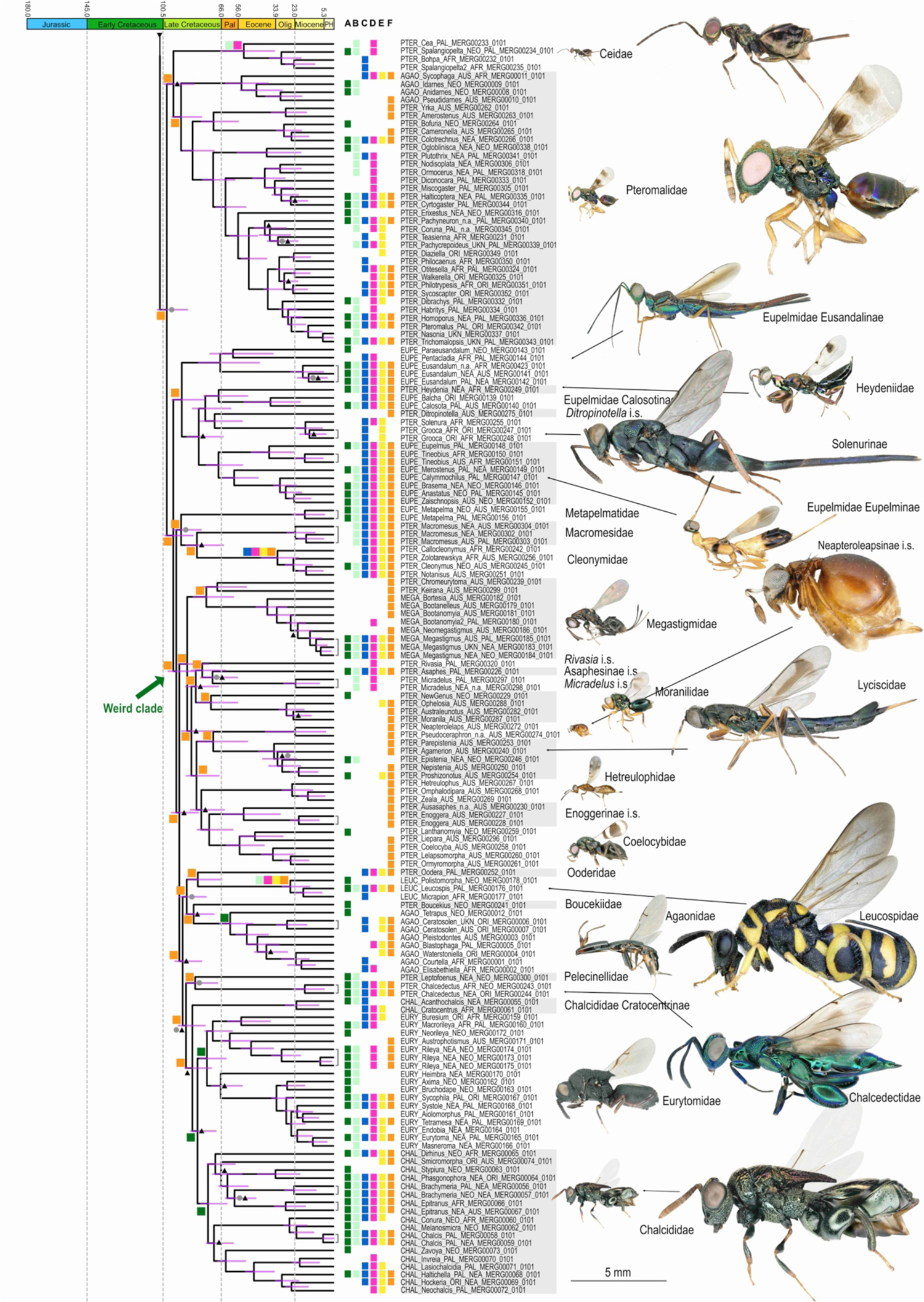
Global historical biogeography of Chalcidoidea and tentative new classification. The chronogram obtained from the complete set of ingroup taxa is illustrated and current classification is used to annotate tips (four letter prefixes; see also Table S1 for complete information on sampling). The tentative new familial classification is materialized with successive grey and white boxes on tip labels and shown next to the tree. For clarity, ancestral ranges (BAYEAREALIKE+*J*) are given only up to family level. The complete scenario and alternative inferences of ancestral ranges are provided in Figure S6. Inferences of ancestral ranges were conducted with only one specimen per genus as shown with brackets that connect tips. Current distribution of genera is shown with colored boxes at tips. Sampling area of specimens is indicated in tip labels. NEO=Neotropical; NEA=Nearctic; AFR=Afrotropical; PAL=Palaearctic; ORI=Oriental; AUS=Australasian. UKN=Unknown when collection data are unavailable. Stars indicate that specimens were sampled in areas where species was introduced or not yet cited. Sampling area for the specimen used for sequencing exons is listed first, sampling area for the specimen used for sequencing UCEs is listed second; n.a. is used when no specimen was sequenced and only one sampling area is reported when exons and UCEs were obtained from specimens sampled in the same areas (or from the same specimen). Unless specified, nodes are supported by SHaLRT > 80%, UFBoot > 95% and sCF > 34.3 (minimum support for a family that is well defined morphologically, Trichogrammatidae). Nodes with a grey circle are supported by SHaLRT < 80% or UFBoot < 95%; nodes with a black circle are supported by SHaLRT < 80% and UFBoot < 95%; nodes with a black triangle are supported with sCF < 34.3. Images on the left of tentative family names are all at the same scale. Images on the right of tentative family names have been magnified. Photos ©K. Bolte (Baeomorphidae); ©J.-Y. Rasplus (all others).

### Divergence time estimates and historical biogeography

The chronogram obtained from the combined tree is shown in Figures 4 and S5. Divergence time estimates and confidence intervals for all nodes are given in Table S3b. Estimates of divergence time indicate that Chalcidoidea diverged from their sister group (Mymarommatoidea) 174.0 million years ago (Ma) [95% Equal-tail Confidence Interval (95% CI) 167.3–180.5 Ma]. Crown Chalcidoidea is dated at 162.2 Ma (153.9–169.8 Ma). The four first splits on the backbone (Mymaridae; Rotoitidae; “Tiny Wasp clade”; all other chalcid wasps) occur over a time span of ∼53 million years (Myr). From ∼ 110 Ma, divergences are closer in time with the remaining 8 splits on the backbone spanning ∼ 24 Myr. Ancestral range estimations for all biogeographical models are provided in Figure S6. AICc favored the BAYAREALIKE+*J* model (Table S4c; Figure 4). A South Gondwanan origin of Chalcidoidea [Australasian (DEC, DEC+*J*, BAYAREALIKE+*J;* DIVALIKE+*J*) or Australasian+Neotropical (BAYAREALIKE; DIVALIKE)] is suggested by all models, with colonization of the rest of the world more or less delayed in time depending on the model used (Figure S6).

## DISCUSSION

### Mutational saturation disturbs phylogenomic inferences

This study expands upon earlier investigations (Munro et al. 2011; Heraty et al. 2013; Peters et al. 2018; Zhang et al. 2020) to yield a more comprehensive phylogenetic framework for higher relationships within Chalcidoidea. We used a comparison of phylogenomic inferences from molecular data sets (exons for 414 taxa and UCEs for 407 taxa) and morphological/biological/ecological knowledge to enable us to detect systematic bias attributable to mutational saturation, and, conversely, morphological convergence. With decreasing saturation, exons and UCEs topologies tended to become more similar (Figures 1-2, S1, Appendix S2) and relationships of groups for which placement could be evaluated based on morphological data tended to become more concordant with morphology or biology (Table S6), though not always. Thus, we confirm that mutational saturation is an important source of error in phylogenomics, especially for deep-time inferences (Borowiec et al. 2015; Borowiec 2019; Duchêne et al. 2022). We comment below on the groups of taxa that revealed inference bias attributable to saturation and/or for which morphology was misleading.

### Mymaridae and Rotoitidae

Mymaridae was recovered as sister to all other Chalcidoidea with strong support in all but the most saturated UCE data set. With this last data set, Rotoitidae was inferred as sister to the rest of chalcid wasps including Mymaridae. With decreasing saturation, the SHaLRT and UFBoot support for Mymaridae as sister to other Chalcidoidea increased in the UCE trees. Mymaridae and Rotoitidae have long been hypothesized as the first and second lineages of Chalcidoidea to diverge from their common ancestor (Gibson and Huber 2000; Munro et al. 2011; Heraty et al. 2013). We provide, for the first time, strong molecular support for this hypothesis.

#### Agaonidae

Sycophaginae and all other subfamilies of Agaonidae are both associated with *Ficus* (Moraceae). They have been considered as part of the same family based on morphological data (Heraty et al. 2013), but this result was not supported by earlier molecular data (Munro et al. 2011). In our results, Sycophaginae consistently clustered away from other Agaonidae. With decreasing saturation in the exon data set and with UCEs, Sycophaginae were consistently recovered in the same group of pteromalids. While several morphological characters group Sycophaginae and other Agaonidae, several others are shared with lineages of Pteromalidae with which Sycophaginae clustered on our trees. These include the separated postgenae with an interceding lower tentorial bridge impressed relative to the postgena, the structure of the antenna 14-segmented with a terminal button, and presence of an axillular sulcus in all but a few highly derived species (Heraty et al. 2013). In addition, the gall-associated biology of Sycophaginae is similar to that of Colotrechninae, another lineage with which Sycophaginae cluster.

#### Chalcididae and Eurytomidae

The polyphyly of Eurytomidae as well as the placement of Cratocentrinae away from other Chalcididae in the most saturated exon and UCE data sets can be also attributable to mutational saturation. Indeed, Eurytomidae is a well-supported family (Heraty et al. 2013). Cratocentrinae is considered as the sister group of other Chalcididae (Cruaud et al. 2021). However, a monophyletic Chalcididae is only recovered in the ASTRAL trees (exonsAA, UCEs90-25, combined), while Cratocentrinae is sister to Eurytomidae+other Chalcididae in the IQTREE trees. A mesothoracic spiracle that is hidden in Eurytomidae and all Chalcididae except for Cratocentrinae gives support to Cratocentrinae being sister to the two other taxa, but 15 synapomorphies support Chalcididae as monophyletic (Cruaud et al. 2021). This suggests that despite high statistical support (SHaLRT=94/UFBoot=97 for exonsAA; 100/100 for UCEs90-25 and combined) IQ-TREE inferences may have been misled in this case, but this result requires more investigation.

#### Calosotinae (Eupelmidae)

Although morphology weakly supports Calosotinae as monophyletic (Gibson 1989; Heraty et al. 2013), it was always recovered as polyphyletic in our analyses. Indeed, a group of Calosotinae (*Eusandalum*, *Pentacladia* and *Paraeusandalum*), which exhibit V-shaped notauli, never clustered with other Calosotinae that show paramedially parallel notauli (Gibson 1989). This group is instead more closely related to several Pteromalidae genera (*Heydenia*, *Ditropinotella*, *Grooca* and *Solenura*), a result that is somewhat corroborated by morphology. With decreasing saturation, species belonging to Calosotinae were less scattered across the trees, and Calosotinae with V-shaped notauli became sister to the clade composed of other Calosotinae, some Pteromalidae genera and Eupelminae, with the result that a core group of Eupelmidae (Calosotinae and Eupelminae) was not monophyletic. This result was similarly supported by all of our preferred phylogenomic data sets (Figure S3). The convergent modification of mesosomal structure (enlarged acropleuron of females) linked to the ability to jump in this group could have misled morphological studies (Peters et al. 2018; Zhang et al. 2020).

Notably, none of the studies based on molecular data alone have supported monophyly of a clade with jumping abilities that includes Eupelmidae, Cynipencyrtidae, Encyrtidae, Tanaostigmatidae and some Aphelinidae. Monophyly of Calosotinae, Eupelmidae and a clade that included Cynipencyrtidae, Tanaostigmatidae and Encyrtidae was only found in the combined morphological and molecular analysis of Heraty et al. (2013). Modifications linked to the ability to jump may be at the origin of one of the two characteristics considered as apomorphies defining the clade (mesoscutal lateral lobes “shoulder-like” on either side of pronotum). The presence of parapsidal lines in Calosotinae and in *Solenura* and *Grooca* (Gibson et al. 1999) is a potential argument to redefine Calosotinae, with not all taxa having an enlarged acropleuron. Whether Calosotinae (or Eupelmidae) were wrongly considered monophyletic based on morphology or whether bias remained in the molecular analysis requires more investigation.

#### Encyrtocephalus

As saturation decreased, *Encyrtocephalus* grouped within the Gall clade. Morphologically, *Encyrtocephalus* shares characters with Melanosomellini and only differs from a majority of them by the large supracoxal flange of propodeum and a curved stigmal vein. However, in all our analyses but one (ASTRAL exonsAA), *Encyrtocephalus* never grouped with Melanosomellini but instead is recovered sister to Tanaostigmatidae, a result that also requires more investigation.

It was not possible to assess whether further attempts in reducing bias in fact reduced undetected bias or simply resulted in loss of phylogenetic signal. In all these attempts, placement of Cratocentrinae, *Tetrapus* and *Encyrtocephalus* remained unchanged. No objective criterion has been proposed so far to determine what fraction of genes/sites should be removed from a data set to converge to a ‘correct’ topology and it is unlikely that such a criterion will emerge in the future. Therefore, we advocate that the IQ-TREE combined tree is the best compromise we could achieve with this data set, current evolutionary models and inference methods. Results may be improved in the future either with increasing taxonomic sampling and/or better evolutionary models, given that analyses are computationally tractable. Notably the increased taxonomic sampling of the AHE520 data set yielded nearly identical results (Figures S1, S3; Appendix S2). However, bias will be difficult to track and alternative relationships hard to evaluate given the versatility of morphological characters in chalcid wasps. Indeed, the astounding diversity of morphologies that evolved in about 160 Myr will continue to complicate the finding of strong synapomorphies to support many of the groups. To our knowledge, no morphological analysis provides convincing evidence to reject the global topology inferred with the molecular data set, but there are data suggesting that alternative placements of a few groups are as plausible as those recovered here.

### Validity of current families

Of the 25 currently recognized (extant) families, seven were recovered as paraphyletic or polyphyletic in the least biased molecular data sets, which confirms rampant morphological convergence within Chalcidoidea. We briefly list below the main changes to current familial classification that will result from this study. The complete revision in agreement with ICZN rules in which 48 families are recognized will be published elsewhere (Burks et al. submitted) and we emphasize that the current paper does not include nomenclatorial acts. To help readers, these new family names are mapped on Figure 4, and stem and crown ages are listed in Table 2.

**Table 2.**
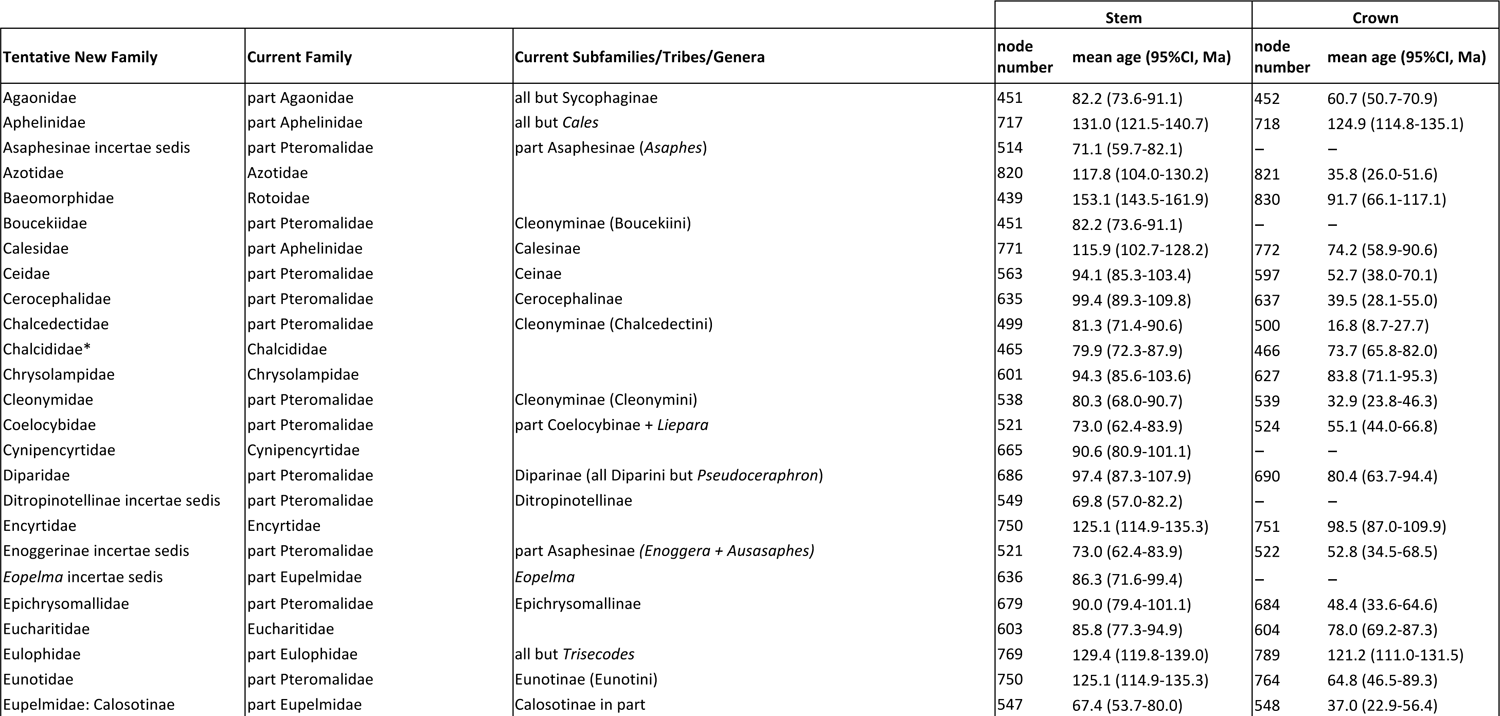
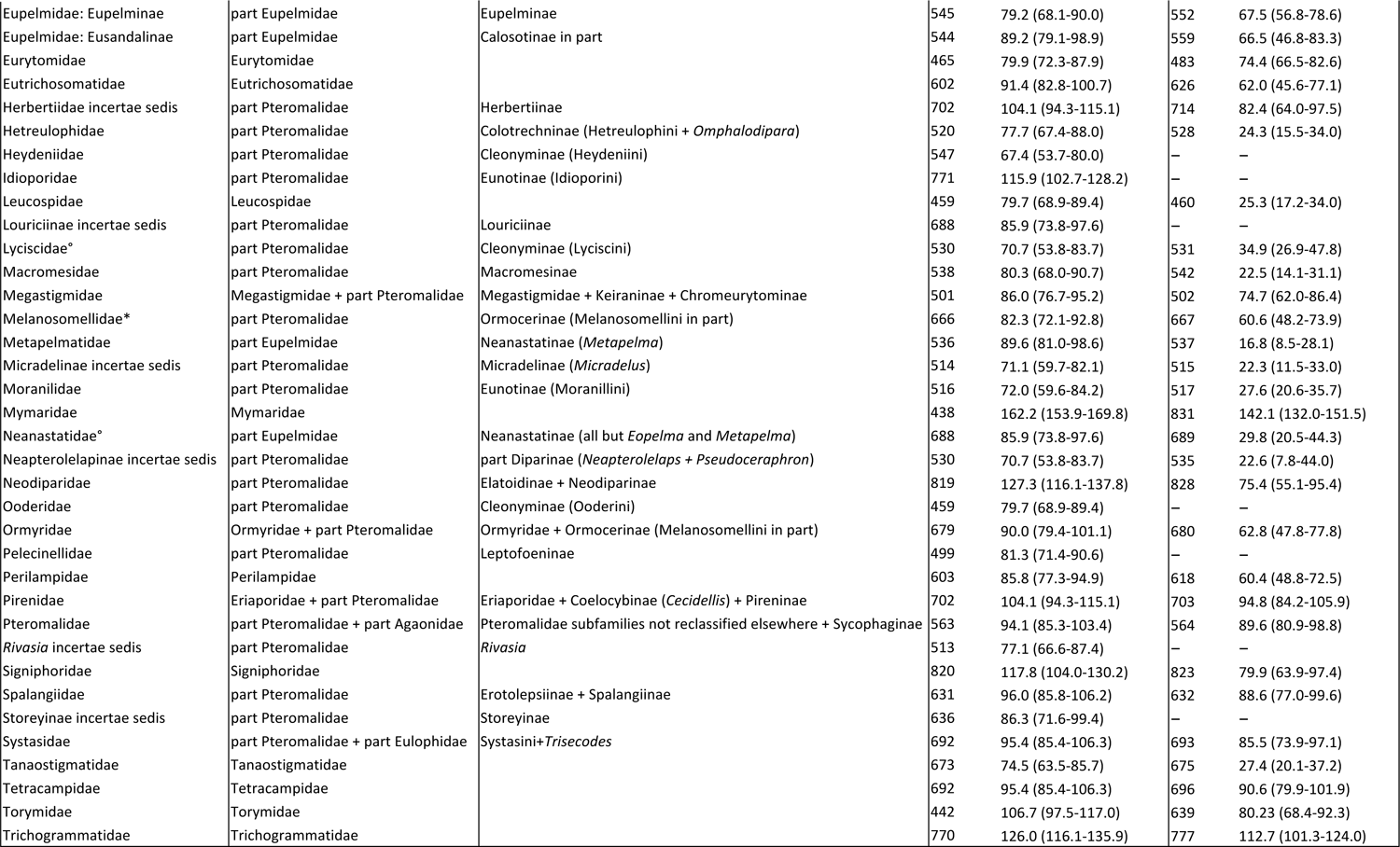
Ages for families of Chalcidoidea. The new classification (Tentative New Family) that is induced by our results will be formally established elsewhere (Burks et al. submitted) but new names are used here for tracking records. Current and tentative new classification of each sample included in our study can be found in Table S1b, see also Figure 4. Node numbers refer to tree in Table S3. Chronogram is available from Figure S5 and Appendix S2. *Ages are given for Chalcididae minus Cratocentrinae; Melasomellidae minus *Encyrtocephalus*; Lyscicidae minus Solenurinae; Neanastatidaae minus *Lambdobregma* but the polyphyly of these clades could be due to artefacts (see text).

#### Aphelinidae

Calesinae clustered away from other aphelinids in all molecular analyses despite morphological affinities (Heraty et al. 2013). We propose that Aphelinidae should be restricted to four subfamilies: Aphelininae, Coccophaginae, Eretmocerinae, Eriaphytinae, while Calesinae should be upgraded to family rank (Burks et al. submitted).

#### Agaonidae

Following discussion in the previous section, Sycophaginae should be removed from Agaonidae.

#### Chalcididae

Until proven otherwise and following the discussion in the previous section, Cratocentrinae is maintained within Chalcididae.

#### Eulophidae

As previously reported with molecular data (Burks et al. 2011; Munro et al. 2011; Heraty et al. 2013; Rasplus et al. 2020), *Trisecodes* (*incertae sedis* within Eulophidae) never clustered with other eulophids, but its placement remains ambiguous. *Trisecodes* is either recovered as sister to Systasini (exonsAA 433/520, combined) or Trichogrammatidae (UCEs90-25). *Trisecodes* exhibits the 3-segmented tarsi of Trichogrammatidae, but this is a characteristic that has occurred several times independently across Chalcidoidea. *Trisecodes* shares with Systasini the presence of a mesofurcal pit on the mesotrochantinal plate between the mesocoxal insertions, which suggests a closer relationship between the two groups. Defining a family grouping for *Trisecodes* and Systasini seems the best solution, even though *Trisecodes* differs from Systasini in tarsomere and flagellomere count (Burks et al. submitted).

#### Eupelmidae

This family was never recovered as monophyletic in our analyses and no single morphological feature is unique to Eupelmidae (Gibson 1989; Heraty et al. 2013), which casts doubt on its validity. In all molecular trees, Neanastatinae (including *Metapelma*) and Calosotinae are polyphyletic while Eupelminae are recovered as monophyletic. Furthermore, *Eopelma* never groups with other eupelmid clades and is instead consistently recovered sister to *Storeya*, the unique genus of Storeyinae (Pteromalidae). Status and placement of the current genera and subfamilies of Eupelmidae are thoroughly discussed elsewhere (Burks et al. submitted).

#### Eriaporidae

In all topologies, *Cecidellis* (Pteromalidae) renders Eriaporidae paraphyletic. However, although the genus is well defined by a short lamina covering the posterior propodeal margin in the female, it resembles some Eriaporidae in coloration and eye divergence, and Pireninae in body shape features. In addition, Eriaporidae is recovered sister to Pireninae (Pteromalidae) in all our reconstructions, a group with which it shares several morphological characters. Therefore, we advocate a single family grouping of *Cecidellis*, Eriaporidae and Pireninae (Burks et al. submitted).

#### Pteromalidae

Pteromalidae *s.l.* is scattered across all inferred trees. This hyperdiverse family that contains 33 subfamilies and nearly 650 genera has long been considered as a “taxonomic garbage can” for taxa that could not be easily included in other chalcid families (Burks et al. submitted). Pteromalidae has long been shown to be polyphyletic (Munro et al. 2011; Heraty et al. 2013; Peters et al. 2018; Zhang et al. 2020), but the lack of robust support for previous phylogenies has precluded any taxonomic rearrangement. Although the backbone of our Chalcidoidea tree remains unresolved in some places, shallower clades are strongly supported that will enable a first revision of Pteromalidae (Burks et al. submitted). Our results do support a new Pteromalidae *s.s*. that includes a number of odd fig-wasp parasitoids and the Sycophaginae that were previously treated as Agaonidae. Twenty families, that mostly correspond to the current subfamilies or tribes of pteromalids apart from Pteromalidae *s.s.*, will need to be erected and their circumscription redefined (Figure 4; (Burks et al. submitted). Most of these new families are ancient lineages (100-80 Ma, Table 2) that were likely grouped together within Pteromalidae based on symplesiomorphies, or because they had no apparent affinities with any other family. Other subfamilies will be included in other existing families that will be redefined (e.g., Keiraninae and Chromeurytominae within Megastigmidae) or will remain *incertae sedis* (e.g., Storeyinae) (Burks et al. submitted).

### Higher level relationships

From here on, family names refer to the new classification that will be formally established elsewhere (Burks et al. submitted); see also Figure 4 and Table 2). Although it is difficult to say whether it is due to a lack of signal or noise creating conflicting signal across the genome as previously suggested (Zhang et al. 2020), relationships in some places are still unresolved. Depending on which measure of statistical support is accounted for, the backbone is either a rake or moderately supported (Table S2e; Figures 3, 4, S1). Only two backbone nodes are recovered in the set of gene trees (#1 and #2; Figures 3, 4), likely because of a lack of signal linked with the short length of loci that results in unresolved gene trees (Figure 3B). Three backbone nodes (#4-6; Figures 3, 4) do not receive any significant statistical support (gCF, sCF, UFBoot and SH-aLRT) and the corresponding part of the topology should be better regarded as a polytomy. As for the early evolution of birds (Suh 2016), phylogenomics may not be able to resolve these difficult nodes because of near-simultaneous speciation. Indeed, from backbone node #2 (“Tiny Wasp clade”), one lineage appears, on average, every 3 Myr (Table S3b, Figure 4) and the branching pattern is characterized by very short branches that likely lead to gene tree incongruence.

Interestingly, certain higher-level relationships that reflect shared biology more than morphology were inferred with decreasing saturation in the individual data sets or emerged in the combined tree in agreement with hidden support in the individual data sets (Figure 3A).

### “Planidial clade” (Eucharitidae + Perilampidae + Eutrichosomatidae + Chrysolampidae)

We confirm that families with planidial larvae (Zhang et al. 2022) form a monophyletic group that is recovered in all trees. Statistical support for this clade in the combined tree is high (SHaLRT=100/UFBoot=100/gCF=0.197/sCF=36.5) with, again, the exception of gCF. Sister to the “planidial clade” we recovered a clade of two old pteromalid lineages that split between 89 and 96 Ma that should be considered as one family (Spalangiidae; Burks et al. submitted). Only larvae of Spalangiinae have been described so far (Tormos et al. 2009) and, interestingly, *Spalangia* larvae appear to be mobile (Gerling and Legner 1968), with a series of tubercles across the ventral region of body segments II–XII (Tormos et al. 2009). This latter feature is also documented in several lineages of the planidial clade [Chrysolampidae: Chrysolampinae (Askew 1980; Darling and Miller 1991); Philomidinae (Darling 1992); and Eutrichosomatidae (Baker and Heraty 2020)], which may corroborate the close relationship of these taxa.

### “Gall clade” (Cynipencyrtidae + Epichrysomallidae + Melanosomellidae + Ormyridae + Tanaostigmatidae)

This higher-level grouping is recovered here for the first time. Again, with the exception of gCF, statistical support in the combined tree is high (100/100/0.049/34.2). On the morphological side, there is limited evidence to support this clade that groups wasp lineages previously classified in four different families. However, from the perspective of life-history strategy all lineages are gall-associated wasps. Melanosomellidae, Epichrysomallidae and Tanaostigmatidae are gall-makers associated with several groups of angiosperms (e.g. *Nothofagus*, *Casuarina*, *Eucalyptus*, *Ficus* and legumes, among others; LaSalle 1987; Bouček 1988; Beardsley and Rasplus 2001; LaSalle 2005). *Asparagobius* is recovered sister to *Ormyrus* and is also a gall-maker on *Asparagus* in Africa. *Ormyrus* has been demonstrated to be parasitoid of gall-makers (Gomez et al. 2017), while *Cynipencyrtus* is a parasitoid of either Cynipidae or their inquilines (Ito and Hijii 2000).

### “Tiny Wasp clade” (Aphelinidae + Azotidae + Calesidae + Neodiparidae + Encyrtidae + Eulophidae + Eunotidae + Idioporidae + Signiphoridae + Trichogrammatidae)

This higher-level grouping is highlighted for the first time but recovered only in the IQ-TREE combined tree with moderate support (100/74/0/30.9). However, there is hidden support for this clade in the exonsAA (sCF=32) and UCEs90-25 (sCF=31.4) data sets (Figure S1). In addition, the backbone node (#3) that splits Mymaridae/Baeomorphidae/”Tiny Wasp clade” from the other Chalcidoidea is moderately supported (100/73/0/37.9) and receives hidden support from the exonsAA (sCF=38.6) and the UCEs90-25 (sCF=40.4) data sets (Figure S1). Statistical support for the clade is possibly affected by the ambiguous placement of the rogue *Trisecodes* (cf. previous section) that is either nested within this clade (UCEs) or that clusters with Systasidae and Tetracampidae (exonsAA). From a morphological point of view, the “Tiny Wasp clade” has hardly any support other than usually being small and mostly soft-bodied (except some Eulophidae). The clade comprises several lineages characterized by a reduction in the number of flagellomeres or tarsomeres, and the frequent presence of a mesophragma that extends into the metasoma through a broad union with the mesosoma. Biologies are diverse in this group and there are no clear trends. With the exception of Eulophidae, that parasitize nearly all insect orders, lineages of the “Tiny Wasp clade” are more frequently associated with Hemiptera. They appear to be mainly endoparasitoids of exophytic hosts (such as mealybugs), while other chalcid wasps are more frequently ectoparasitoids of endophytic hosts. In the same manner, species of this clade also appear to be more frequently oophagous than other chalcid wasps. Confirmation of this clade as a monophyletic lineage requires increasing taxonomic sampling, which may help to stabilize the placement of *Trisecodes* as well as a formal study of the evolution of host-associations within Chalcidoidea.

### “Weird Clade” (Megastigmidae + Leucospidae + Agaonidae + Metapelmatidae + Eurytomidae + Chalcididae + several poorly diversified families)

This higher-level grouping was unexpected based on morphology or biology. Yet, this “Weird clade” is statistically slightly better supported than the “Tiny Wasp clade” in the combined tree (99.2/95/0/35.6). Hidden sCF support for this clade is 35.9 for UCEs and 34.3 for exonsAA. From a morphological point of view the “Weird clade”, as its name indicates, groups disparate lineages of chalcid wasps that exhibit contrasting morphologies which could be correlated with their diverse biologies (Figures 3 and 4). Many families such as Boucekiidae, Chalcedectidae, Cleonymidae, Lyciscidae, Metapelmatidae, and Pelecinellidae, but also several genera of Eurytomidae and a few Chalcididae are parasitoids of xylophagous insects (mostly Coleoptera). Leucospidae have shifted to solitary bees and wasps that nest in wood or in mud nests, but *L. dorsigera* has also been reported as a hyperparasitoid of xylophagous beetles (Hesami et al. 2005). Agaonidae, Megastigmidae and Eurytomidae are mostly phytophagous, but the last two families also comprise parasitoid species. Agaonidae enter figs (inflorescences of *Ficus*, Moraceae) through a small aperture called the ostiole and exhibit strong morphological adaptations (mandibular appendage, anelli fused into a hook-like process, short protibia with spurs, etc.; Cruaud et al. 2010). Some lineages have more specialized biology such as Moranilidae that are predators of mealybug eggs, Asaphesinae (*incertae sedis*) that are hyperparasitoids of aphids through Braconidae wasps, and Enoggerinae (*incertae sedis*) that are oophagous parasitoids of Coleoptera. Coelocybidae are gall-associated wasps. Finally, Chalcididae attack nearly all insect orders, but mostly Lepidoptera. From our collective knowledge, it is hard to determine whether this “Weird clade” results from an inference bias or accurately reflects phylogenetic affinities.

Morphology is rather uninformative in its support for most higher-level relationships. Nevertheless, we find a dichotomy between early diverging lineages that are small “soft-bodied” chalcid wasps prone to shriveling when air-dried (Mymaridae, Baeomorphidae, “Tiny Wasp clade”, Pirenidae) and “hard-bodied” chalcid wasps that diversified later (Figures 3,4). Higher level morphological convergences are confirmed as exemplified by the evolution of an enlarged acropleuron and correlative transformation of legs linked to the ability to jump that may have happened at least 7 times independently during the evolutionary history of chalcid wasps (in 1) Encyrtidae sister to Eunotinae, 2) *Lambrodegma* and *Neanastatus* sister to *Callimomoides*, 3) *Eopelma* sister to *Storeya*, 4) *Metapelma* sister to *Macromesus*+Cleonymidae, 5) Tanaostigmatidae sister to *Encyrtocephalus*, 6) *Cynipencyrtus* sister to Melanosomellidae + the preceding clade, 7) Calosotinae sister to *Heydenia*; all relationships excepted *Metapelma* with maximum support).

Finally, with the exception of the exonsAA and AHE520AA trees (low support for Diapriidae as sister to Chalcidoidea), all of our results strongly support the sister group relationship between Mymarommatoidea and Chalcidoidea. Gibson (1986) was the first to propose a sister group relationship between these two superfamilies. Munro et al. (2011) recovered Diaprioidea+Mymarommatoidea as sister to Chalcidoidea, although when combined with morphology (Heraty et al. 2013), mymarommatids were sister to Chalcidoidea.

### Time line and historical biogeography

An important result of our study is a revision of the temporal scale over which Chalcidoidea have evolved and dispersed throughout the world (Figures 4, S5, Table S3). Likely because our taxonomic sampling is one or two orders of magnitude higher than previous phylogenomic studies (Branstetter et al. 2017; Peters et al. 2017; Peters et al. 2018; Tang et al. 2019), we infer an older crown age for Chalcidoidea: 162.2 (154.0-170.0) Ma. Nevertheless, this age falls within the confidence interval of the only other Chalcidoidea-centered time tree [129Ma (89-208); (Peters et al. 2018)]. Importantly, our estimates are compatible with the meager fossil records of chalcid wasps but also with Earth’s paleo-geological history (AppendixS1; Figure 4).

The oldest fossils of chalcid wasps belong to the so-called “soft-bodied” chalcid wasps (Mymaridae, Baeomorphidae and “Tiny Wasp clade”) (Haas et al. 2018). The oldest putative chalcid fossil is *Minutoma yathribi* Kaddumi, 2005 from Jordanian amber (Albian, ∼113.0– 100.5 Ma) (Kaddumi 2005). But the uncertain affinities of this fossil preclude us to use it in our analyses. The oldest unambiguous chalcid fossils are a Mymaridae [*Myanmymar aresconoi*des; Poinar & Huber (2011)] and a Baeomorphidae (*Baeomorpha liorum*; (Huber et al. 2019) from Myanmar (Burmese) amber (minimum age 98.2 Ma; Appendix S1). Species of Baeomorphidae are also frequent in Cretaceous ambers of the northern-hemisphere [in the retinites of Baikura (minimum age 94.3 Ma) and of Yantardakh (minimum age 83.6 Ma; Gumovsky et al. 2018) and Canadian ambers of Cedar Lake and Grassy Lake, which are Campanian in age (83.6-72.1 Ma; McKellar et al. 2008)].

Only a dozen fossils of “hard-bodied” Chalcidoidea are known from Cretaceous formations. Among them, three were not used as calibrations because they had uncertain affinities which prevent them from being assigned to a clade. Nevertheless, all of them fit relatively well within the proposed time-frame for Chalcidoidea. Diversinitidae (Myanmar amber) possibly belongs to the group of “hard bodied” taxa (Haas et al. 2018), however it has uncertain relationships to extant chalcid wasps. Two other undescribed fossils clearly belong to this large clade: 1) a few specimens with uncertain morphological affinities (Pirenidae or Micradelinae) (A. Gumovsky and M.-D. Mitroiu, pers. comm.) from Taimyr amber (86.3-83.6 Ma), and 2) an unidentified “torymid” specimen from Canadian amber (83.6-72.1 Ma) (Figure 4C in McKellar and Engel 2012) that probably belongs instead to an extinct lineage.

Splits between Neotropical and Australian lineages also corroborated our new time frame for chalcid wasps. Indeed, several clades whose ancestor probably dispersed through the Antarctic land bridge predate connection break-ups and temperature decreases (i.e., 45 Ma; van den Ende et al. 2017). Thus, the clade grouping the Australian *Aeschylia* with the Neotropical *Aditrochus* and *Plastobelyta* (Melanosomellidae), all of which are gallers on *Nothofagus*, is dated at 60.1 Ma (Figure S5, Table S3b), while the split between the Australian *Liepara* and the south Andean *Lanthanomyia* (Coelocybidae) that is, among others, a parasite of *Aditrochus* species is dated at 46.2 Ma. The split between the Neotropical *Erotolepsia* and its Australian/Oriental sister *Papuopsia* (Spalangiidae) is estimated at 49.9 Ma. The stem age of neotropical Lyciscidae that are nested within an Australian clade is estimated at 33 Ma, which is a very late time to cross Antarctica.

Only a few dating analyses have been performed for groups of chalcid wasps. Variability in mean age estimates were noted depending on the study, data sets and methods used (differences up to 12 Myr). Our estimates for the age of the planidial clade [94.3 Ma (85.6– 103.6)] and Eucharitidae [78.0 Ma (69.2–87.4)] are close to previous estimates (Murray et al. 2013; Zhang et al. 2022). The mean age for crown Agaonidae at 60.7 Ma (50.7–70.9) is 15 Myr younger than previous estimates of 75 Ma (94.9–56.2) (Cruaud et al. 2012) but this difference is likely due to the reduced set of taxa used in our study. Nevertheless, ages of all other groups of wasps that are strictly associated with *Ficus* (Epichrysomallidae, Sycophaginae, and other pteromaline fig wasps) postdate the age of Agaonidae.

The origin of Chalcidoidea and divergence of the two first lineages (Baeomorphidae 153.1 Ma; “Tiny Wasp clade” 136.1 Ma) coincide with a rise in insect fossil families in the late Jurassic to the Hauterivian–Barremian (Schachat et al. 2019). The next splits on the backbone (from 110.3 Ma) coincide with the second sharp increase in fossil diversity through the Albian and Cenomanian (Schachat et al. 2019). This rapid radiation of “hard-bodied” Chalcidoidea, between 110 and 80 Ma coincides with the onset and diversification of flowering plants and holometabolan insects. Although this hypothesis should be formally tested through a thorough compilation of host associations and reconstruction of ancestral life histories, the first lineages to diverge (Mymaridae, Baeomorphidae, “Tiny Wasp clade”) are first likely oophagous and subsequently mostly associated with Hemipteran hosts (Aphelinidae, Azotidae, Calesidae, Encyrtidae, Eunotidae, Signiphoridae). Recently it was discovered that the sister group to Chalcidoidea, Mymarommatoidea, are parasitoids of the eggs of Lepidopsocidae (Psocoptera) (Honsberger et al. 2022), thus adding further support to an ancestral habit of egg parasitism for Chalcidoidea. Subsequently, chalcid wasps switched to virtually all orders and life stages of Holometabola. Interestingly, the first shifts to phytophagy (e.g., stem-crown gall clade: 102.1-98.2 Ma) correspond to the beginning of the “Angiosperm Terrestrial Revolution” ca. 100 Ma (Benton et al. 2022).

We also propose a biogeographical scenario for Chalcidoidea that should be regarded as a first hypothesis. Indeed, although our sampling is highly representative of the main lineages occurring on Earth (very few suprageneric taxa are missing) and representative of the distribution of described genera (Noyes 2019), several hyperdiverse clades have been scarcely sampled (Trichogrammatidae, Encyrtidae, etc.). Furthermore, the analyses could be driven by our sampling of endemic genera that is slightly biased compared to the overall diversity of endemic chalcid genera. Finally, our knowledge of the current diversity of genera and species in several biogeographic regions remains limited and future discoveries may change some of our inferences.

Our most likely biogeographic scenario (BAYAREALIKE+*J*) inferred an East Gondwanan (Australia) origin for Chalcidoidea. It also suggested that the Cretaceous and rapid radiation of Chalcidoidea occurred in southern Gondwana. This scenario favored a ‘multiple dispersal out of Australia’ hypothesis, with the last dispersal event on the backbone being the ancestor of Eurytomidae and Chalcididae, ca. 83 Ma, ca. 28 Myr before the disconnection of South America and Antarctica (Reguero et al. 2014). All other scenarios confirm a south Gondwanan origin in Australasian (DEC, DEC+*J*, DIVALIKE+*J*) or in Australasian+Neotropical areas (BAYERALIKE and DIVALIKE).

While there is no formal biogeographic analysis of Mymaridae, their East-Gondwanan origin is reasonable. Indeed, the Australian fauna contains the earliest diverging lineages of Mymaridae as well as a large diversity of genera that encompasses all familial subgroups found in the family (Lin et al. 2007).

Extant genera of Baeomorphidae occur in southern temperate rainforests of Chile (*Chiloe*) and New Zealand (*Rotoita*) and show a disjunct amphi-Pacific distribution characteristic of taxa that once ranged in southern Gondwana across Antarctica (van den Ende et al. 2017). The split between these genera (91.7 Ma) appears contemporaneous with the breakup of southeast Gondwana and with the drift of Zealandia away from Antarctica (between 95 and 84 Ma) (Schellart et al. 2006; Mortimer et al. 2017).

Recent findings by J.M. Heraty and C. Dominguez (pers. comm.), strongly suggest that Baeomorphidae may be egg parasitoids of Peloridiidae (moss-feeding bugs) and/or of Myerslopiidae (tree-hoppers), two ancient families of Hemiptera, that are respectively sister to all other Auchenorrhyncha and to all other Membracoidea (Johnson et al. 2018).

Interestingly, both peloridiids and myerslopiids also have an amphi-pacific distribution and Myerslopiidae fossils were only found at Crato in Brazil (122.5–112.6 Ma). In addition, the split between peloridiids from Chile and New Zealand [98 Ma (46–155 Ma); Ye et al. 2019] is close to our estimate for Baeomorphidae and corroborates our time-scale for Chalcidoidea. The fossil record shows that Baeomorphidae were largely distributed in Laurasia, but also on the Myanmar terrane, between the lower Cenomanian and the Campanian (98.2–72.1 Ma) (Gumovsky et al. 2018; Huber et al. 2019), which suggests, in the framework of our scenario, a long northward dispersal. Our results contradict the hypothesis of a Laurasian origin for Baeomorphidae (Gumovsky et al. 2018) that has been made outside of a formal phylogenetic framework. However, given the area of origin for Mymaridae, and the current distribution of *Rotoita* and *Chiloe*, whatever the position of *Baeomorpha*+*Taimyromorpha* (fossil taxa) within Baeomorphidae, it would lead to a South Gondwanan origin for Baeomorphidae and a northward dispersal between 150 and 100 Ma. Their distribution fits well with that of fossils of Coleorrhyncha, the lineage to which Peloridiidae belongs and is now the only extant representative. Coleorrhyncha are divided into two large groups: the Progonocimicidoidea and the Peloridioidea. All extinct families of Peloridioidea occurred in Laurasia (between 201.6 and 112.6 Ma). The oldest fossil of Progonocimicidae occurred in Australia and was dated back to Changhsingian, the uppermost stage of Permian (254.1– 251.9 Ma). Progonocimicidae are recorded from Gondwanan and Laurasian Triassic formations (Evans 1956) but became mostly confined to Laurasia later on (Jurassic and Cretaceous) with the notable exceptions of two Gondwanan fossils found in Cretaceous ambers (Lebanese and Myanmar) (Szwedo et al. 2011; Jiang et al. 2019). Most Coleorrhyncha groups declined during the mid-Cretaceous biotic crisis when vegetation was replaced by modern angiosperms and did not survive the Chicxulub impact (Dong et al. 2014). Our scenario suggests that baeomorphids experienced the same extinction event.

The widespread distribution of most groups included in the “Tiny Wasp clade” makes corroboration of a southern Gondwana origin difficult. Diversity should not be considered as a clue for origin. However, the diversity and placement of some Australian or Neotropical endemic taxa in the topology support a southern Gondwana origin (Figure S6). Within Eulophidae, *Perthiola* (mainly Australian) and *Ophelimus* (Australian) are sister to all Entiinae, while *Aleuroctonus* and *Dasyomphale* (Neotropical) are the first lineages to diverge within Euderomphalini. In addition, although there are not included in our phylogeny, *Cales* is subdivided in two species groups, one occurring mostly in Australasia, the other in the Neotropics (Mottern et al. 2011; Mottern and Heraty 2014; Polaszek et al. 2015).

The Australian origin of the “hard-bodied” chalcid wasps is well supported by 1) the cosmopolitan lineages that have their early diverging taxa occurring in Australia, such as Eucharitidae with Akapalinae (Murray et al. 2013), Perilampidae with *Euperilampus* (Zhang et al. 2022), Pteromalidae s.s. with Sycophaginae (Cruaud et al. 2011), and Colotrechninae as well as Megastigmidae with Keiraninae and Chromeurytominae; 2) lineages that originated in southern Gondwana and are mostly Australian and Neotropical. For example, Melanosomellidae, Lyciscidae, Coelocybidae are diverse in Australia and colonized the Neotropics. Additionally, two other families, Boucekiidae (represented by *Boucekius*) and Pelecinellidae (*Leptofoenus*) could be included here as they have Australian representatives (*Chalcidiscelis* and *Doddifoenus* respectively), that, unfortunately, could not be included in our sampling.

Finally, our analyses identified several subfamilial or familial lineages that colonized the northern hemisphere (Palaearctic or Oriental region) during the Cretaceous, at a time when intercontinental dispersals were difficult. Four of them may derive from ancient dispersal events between Antarctica+Australia and Africa (where they also occur) or from a colonization of the Neotropics followed by dispersal to Africa and recent colonization of the Palaearctic region: Eunotidae between 125 and 65 Ma; the ancestor of Neodiparidae + Signiphoridae + Azotidae between 133 and 127 Ma; Ceidae between 102 and 53 Ma and Cleonymidae between 80 and 33 Ma. Two other apparent long-distance dispersal events are difficult to explain as no members of these clades are presently known outside the Palaearctic region (*Rivasia* and *Micradelus* between 81 and 77 Ma and *Cynipencyrtus* at 90.6 Ma). Some other lineages possibly colonized the northern hemisphere through the Neotropics (*Exolabrum* and *Herbertia* between 104 and 92 Ma; *Trisecodes* between 86 and 41 Ma).

Beyond these lineages, Torymidae was hypothesized to have originated in the Palaearctic region (Janšta et al. 2018). Our analysis strongly suggests a dispersal from Southern Gondwana to Laurasia between 106 and 80 Ma, possibly through Africa. In this case, the first lineages of the family have disappeared there or have not yet been sampled. Another possibility, that may also be the route used by *Eopelma* (ca 86.6 Ma) and a new subfamily close to Erotolepsiinae (ca 96 Ma) to reach the Northern Hemisphere (Sunda), would be through drifting India or through ancient terranes (Hall 2012) or the Trans-Tethyan island arc that separated from northern Australia ca 120 Ma (Westerweel et al. 2019), bringing away the Gondwanan fauna (Poinar 2018).

## CONCLUSION

Chalcidoidea may represent one of the largest radiations of any insect group. The superfamily has deep origins in the Middle Jurassic (95%CI for stem age = 167.3–180.5 Ma) followed by a tremendous diversification in the Paleogene concomitant with the radiation of plants and the insects that feed upon them. We have assembled the largest phylogenomic data set ever to address relationships within 356 genera based on 1007 exons and 1048 UCE loci. By reducing saturation, the results of the independent analyses converged in clade support and also became more congruent with our morphological support for certain family group relationships. Conversely, morphological convergences are highlighted. The combined analysis of the exonsAA and UCEs90-25 data sets (2054 loci and 284,106 sites) produced a generally well supported hypothesis. In some cases, we found clades that matched more with natural history over their morphological support and in others we found some historical morphological groups (i.e., Eupelmidae, Cleonyminae) that were never recovered in our phylogenomic results, which confirm previous analyses (Heraty et al. 2013). In either case, we need to further explore these groups to find the basis of disagreement. Notably, we found a clade of gall-associated wasps that was not previously envisioned, but is indeed a good example of molecules suggesting a clade that is agreeable with re-examined morphology. Our biogeographical inference hypothesizes a general dispersal of taxa from a southern Gondwanan origin. The K-T meteorite event certainly had an impact on the present-day distribution and diversity of chalcid wasps with possible subsequent recolonizations. However, a much larger sampling is required to assess it. Importantly, even with large and independent data sets, the importance of taxonomic evaluation and attempts to reduce saturation and homoplasy were all important factors in developing a concrete phylogenetic hypothesis for this massive radiation. The results support a reclassification of many different chalcid families that will be published elsewhere (Burks et al. submitted), which will set the stage for a better foundation for evaluation of life history attributes across the superfamily.

## ACKNOWLEDGEMENTS

JYR and AC are grateful to Audrey Weber (INRAE, France) for sequencing of the UCE libraries and to the Genotoul bioinformatics platform Toulouse Midi-Pyrénées, France for providing computing resources. We thank Gary Gibson (Agriculture and Agri-Food, Canada), Paul Hanson (Univ. de Costa Rica), Christopher Darling (Univ. of Toronto, Canada), Nicole Fisher (CSIRO, Australia), Michael Gates (USDA, USA), Michael Haas (Univ. of Marburg, Germany), Christer Hansson (Museum of Biology, Sweden), Jason Mottern (USDA, USA), John D. Pinto (UCR, USA), Stefan Schmidt (ZSM, Germany), Christine Lambkin, Chris Burwell and Susan Wright (QM, Australia) for providing specimens and for helpful discussion. AC and JYR acknowledge the Queensland parks and wildlife services for collecting permits (WITK18278817-WIF418664617). We dedicate this work to the memory of our dear friend and colleague John LaSalle, specialist of Eulophidae who was an enthusiastic member of this project.

## DATA ACCESSIBILILTY

Raw paired reads were uploaded as NCBI Sequence Read Archives (PRXXXXX for AHE and PRXXXX for UCEs). Data matrices are available on DRYAD (XXXX). Supplementary data are available upon request from the corresponding authors.

## AUTHOR’S CONTRIBUTION

Designed the study: JYR, JBW, JMH. Obtained funding: JYR, AC, MY, JBW, JMH.

Contributed samples or sequences: JYR, AC, RB, GD, LF, AG, JTH, PJ, MDM, JSN, SVN, AB, JB, HB, BBB, SGB, KB, RSC, NDS, ADM, CD, MG, EG, RLK, LK, EM, JLNA, RKP, RSP, AP, JT, ST, ET, JBW, JMH.

Identified samples: JYR, RB, GD, LF, AG, JTH, PJ, MDM, JSN, SVN, AB, HB, ADM, JLNA, RKP, AP, JT, ST, ET, JBW, JMH.

Organized meetings for sharing knowledge: JYR, PJ, KB, NDS, JBW, JMH.

Performed laboratory work: AC, JYR, BBB, LF, ARL, AML, SN, LS.

Analyzed data: AC, JYR (all but AHE520); JZ (AHE520).

Contributed scripts: AC, JZ, MC, MY.

Compilation of geographical occurrences: JYR, JSN, AC.

Contributed morphological, biological and biogeographical knowledge: JYR, RB, GD, LF, AG, JTH, PJ, MDM, JSN, SVN, AP, JMW, JMH.

Discussed results: JYR, AC, JZ, RB, JBW, JMH.

Drafted the manuscript: JYR, AC, JMH.

All authors revised and commented drafts at different stages and contributed to the final version of the manuscript.

